# Glucose and Oxygen Metabolism Coordinate Human Cortical Developmental Decisions

**DOI:** 10.64898/2025.12.02.691634

**Authors:** Taylor R. Pennington, Alexandria S. M. Morales, Ammar Mehdi, Sophia A. Cerna, Gradi Bamfonga, Jinhua Chi, Alicia M. Saarinen, Sierra Wilferd, Fabiha Firoz, Tran Nguyen, Lingjun Li, Haiwei Gu, Benjamin Bartelle, Christopher Plaisier, Clifford Folmes, Madeline G. Andrews

**Affiliations:** School of Biological & Health Systems Engineering, Arizona State University, Tempe, AZ 85282; Biomedical Engineering PhD Program, Arizona State University, Tempe, AZ 85282; Department of Cardiovascular Medicine, Mayo Clinic Arizona, Scottsdale, AZ 85259; College of Health Solutions, Arizona State University, Phoenix, AZ 85004

## Abstract

Defining metabolic regulation of neurodevelopmental programs is essential to approach developmental disorders and injuries driven by alterations in metabolism. *In vitro* cultures are the only available method to temporally perturb and study living human brain cells throughout neurogenesis, however most culture systems use supraphysiologic conditions of essential nutrients, glucose and oxygen. We probed how environmental exposure to endogenous-like concentrations impact metabolic state and developmental progression of cortical cell types using organoids. Nutrient accessibility globally impacted metabolic state, yet developmental responses to metabolic changes were cell type-specific. Metabolomic and transcriptomic datasets reveal increased TCA metabolites and amino acids and oxidative phosphorylation (OXPHOS) genes, under physiologic glucose conditions. Oxygen level had a modest, yet specific, molecular impact on deep layer excitatory neurons. We assessed consequences of metabolic changes on fate and observed that physiologic glucose expanded the human-enriched population of cortical stem cells, outer radial glia, and their progeny, upper layer excitatory neurons. Alterations in oxygen, instead, affected production of neurogenic progenitors and neuronal differentiation, with higher oxygen availability supporting shifts toward mitochondrial metabolism necessary for maturing cell types. We functionally tested this transition by inhibiting glycolysis; total inhibition promoted neuronal differentiation, whereas inhibition of anaerobic glycolysis/lactate production led to oRG expansion. Lactate signaling was sufficient to suppress oRG development and promote self-renewal of neurogenic progenitors. These data suggest that refined metabolic switches and decreased reliance on glucose are required for transition from stem cell self-renewal to more mature, diversified progenitor subtypes, where switch from anaerobic to aerobic metabolism discretely impacts progenitor diversification.

## Introduction

During embryogenesis, nutrient accessibility within the developing tissue niche regulates bioenergetic activity impacting cellular growth. Yet, the relationship between metabolism and tissue formation has not been fully elucidated. The expansion and organization of the developing brain requires precise coordination of division, differentiation, and fate determination of heterogeneous cell types ^1,2^. Gene expression programs coordinating cellular lineage trajectories have become increasingly revealed due to atlasing strategies ^3–7^. However, the environmental conditions and metabolic states coordinating molecular mediators of cell fate during development are less understood. While functional metabolic analyses have begun in mouse corticogenesis ^8^, the increased complexity of human cortical development is challenging to interrogate. The human cerebral cortex is characterized by protracted cellular expansion and diversification. The bioenergetic needs of increased morphologically and functionally unique populations is not well understood. Clinically, excess or insufficient environmental glucose and oxygen, during gestation or premature birth, as well as rare disorders with gene expression changes in metabolic regulators, are associated with cortical malformations affecting brain size, onset of seizures, intellectual disability, autism, and ADHD ^9–14^. Clinical observations support the relationship between energy metabolism and brain development, but the mechanisms in which metabolic programs regulate developmental trajectories remains unclear.

The availability and influx of bioenergetic substrates, particularly glucose and oxygen, play essential roles in regulating the metabolic state of human cortical cells. Neural stem cells, called radial glia (RG), rely heavily on anaerobic glycolysis to rapidly generate anabolic precursors needed for growth and proliferation ^15,16^. In mice, the anaerobic glycolysis byproduct, lactate, is produced in the progenitor niche, called the ventricular zone (VZ), during early proliferation and lactate inhibition decreases RG proliferation ^8^. During neurogenesis, RG diversify to produce basal progenitors (BPs). In rodents, these are primarily the neurogenic intermediate progenitor cells (IPCs) that subsequently differentiate into the excitatory neurons (EN). Temporal fate decisions result in metabolic shifts to support the energy demands of functionally distinct populations. As tissue growth diminishes, mitochondrial oxidative (OXPHOS) metabolism becomes more prevalent to ensure sufficient energy production for specialized functions of mature cells ^17^. OXPHOS in neurons supports more efficient ATP production to enable signaling activities ^18^ and maturation ^19^. The shift from RG self-renewal to post-mitotic differentiation may accompany transient metabolic shifts throughout differentiation, but the details of this regulation and impact are unclear.

Inappropriate embryonic concentrations of glucose and oxygen lead to neurodevelopmental changes. RG have specific vulnerability to glucose disturbances impacting proliferation, survival, and morphology leading to changes in cortical organization ^20–22^. Adherent human NPC and neuron cultures support that glycolysis genes are required for neuronal survival^23^. Thus, altered glucose utilization elicits consequences for both progenitors and neurons, but the precise impact on developmental dynamics remains unclear. The shift between progenitor expansion to neurogenic differentiation occurs concurrently with angiogenesis ^8^. In addition to cerebrospinal fluid (CSF), the developing vasculature serves as a source of both oxygen and glucose ^8,24^. O_2_ availability is essential for mitochondrial metabolism. RG differentiate into IPCs in sites of angiogenesis ^25,26^, suggesting that oxygen promotes differentiation. However, numerous studies have shown complex, mixed effects of hypoxic states on RG and IPC proliferation and differentiation, rendering its precise impact unclear ^27,28^. Thus, altered quantities of oxygen or glucose during corticogenesis impacts progenitor and neuronal ratios, cortical thickness, and functions with potential life-long impact.

Recently, the source and developmental consequence of metabolic changes in regionalized forebrain organoids has been an outstanding question. While glycolytic gene expression is increased ^29^, the impact of metabolic state has not revealed major neurodevelopmental changes ^30,31^. Gene expression has been the key molecular measure, yet rapid metabolic changes also require functional analysis. Here, we use organoids as a reductionist model of human brain development to directly perturb glucose and oxygen requirement for metabolic state and developmental trajectory. We use orthogonal approaches to temporally assess metabolites, gene expression, and protein-level changes and identify regulation of distinct progenitor subtypes. We observe that glucose regulates outer (o)RG and oxygen regulates IPC production; progenitor shifts lead to alterations in respective neuronal progeny. Temporal fate decisions are associated with shifts in metabolic profile and changes in mitochondrial metabolism. Functional manipulations of glycolysis demonstrate that decreased anaerobic glycolysis and lactate inhibition is required for oRG expansion, while global glycolysis suppression promotes differentiation. Therefore, discrete metabolic programs, guided by nutrient availability, are key for corticogenesis.

## Results

### Culture conditions limit study of metabolic dynamics

Induced pluripotent stem cell (iPSC)-derived 3D organoids are tractable models to evaluate temporal changes in living human cells throughout neurogenesis that can serve as platforms to study gestational injury and developmental disorders ^30^. We leveraged previous temporal characterization of organoid development from human progenitor proliferation (week 4 - 6), diversification (week 6 - 8), and neuronal differentiation (week 10 - 12) of a standard regionalized forebrain organoid protocol ^29,32,33^. Starting from pluripotency (iPSCs) until early gliogenesis (week 16), we did not observe anticipated changes that indicate temporal transitions from glycolysis to cellular respiration via gene expression, metabolite concentrations in spent media, or glucose uptake via fluorescently-labeled glucose analog, 2-NBDG (S Figure 1A-C). Standard forebrain organoids did not reflect anticipated temporal metabolic dynamics predicted from mouse corticogenesis, motivating interrogation of metabolic mediators ^8,25^.

**Figure 1:**
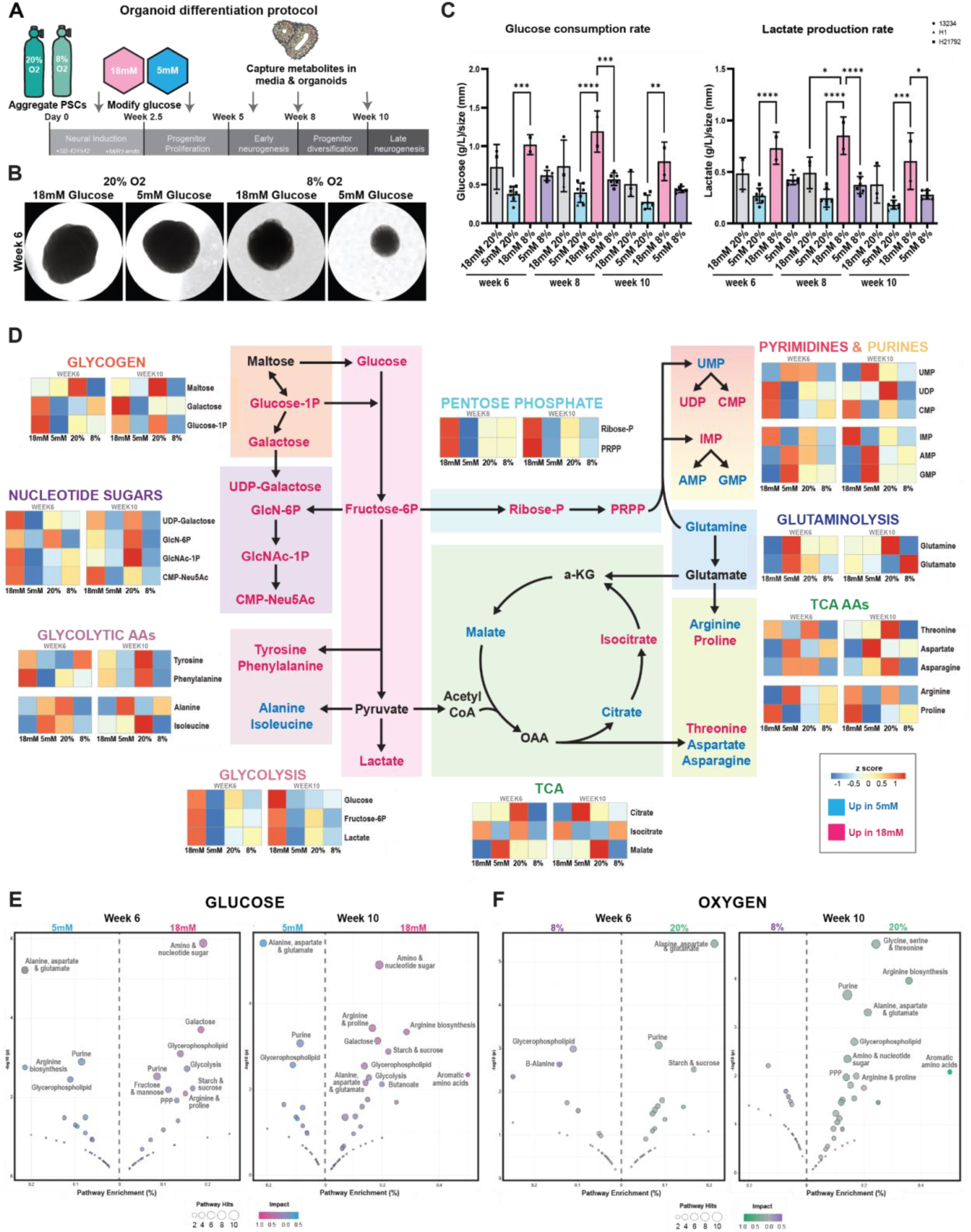
Physiologic glucose and oxygen alter organoid growth rate, metabolic flux, and pathway activity. **A)** Forebrain organoids are aggregated in 8 or 20% O_2_. Glucose is steadily decreased to 5mM concentration from days 9-18. **B)** Brightfield images of week 6 organoids cultured under 20 vs 8% O_2_ and 18mM vs 5mM glucose conditions. **_C)_** Temporal glucose consumption rates per conditions with most significant increases in 8% O_2_ (Ordinary one-way ANOVA with Tukey’s multiple comparisons: Week 6: 5mM 20% vs 18mM 8% ****p<0.0001; Week 8: 5mM 20% vs 18mM 8% ****p<0.0001; Week 8: 18mM 8% vs 5mM 8%: ***p<0.0007; Week 10: 5mM 20% vs 18mM 8%: **<0.0064). Temporal lactate production rates per conditions with most significant increase in 8% O_2_ (Ordinary one-way ANOVA with Tukey’s multiple comparisons: Week 6: 5mM 20% vs 18mM 8%: ****p<0.0001; Week 8: 18mM 20% vs 18mM 8%: *p<0.0279; 5mM 20% vs 18mM 8%: ****p<0.0001; 18mM 8% vs 5mM 8%: ****p<0.0001; Week 10: 5mM 20% vs 18mM 8%: ***p<0.0004; 18mM 8% vs 5mM 8%: *p<0.0251). Datapoints are metabolite concentration values/line ± SD from n>3 spent media collections per timepoint. **D)** Organoids were collected at weeks 6 and 10 for LC-MS. For each timepoint, replicates within each experimental condition were combined. Heatmaps show z-scores comparing metabolites between 18mM vs 5mM and 20% vs 8% O2 for depicted pathways Metabolites were collected from n>5 organoids/timepoint/PSC lines from n=2-3 PSC lines/condition. **E)** Results from MetaboAnalyst Pathway analysis at week 6 and week 10 depict increased pathways in 18mM vs 5mM glucose. Similar observations occurred at both time points. **F)** Results from pathway analysis depict increased pathways in 20% vs 8% O_2_ at weeks 6 and 10. Most pathways were downregulated in 8% O_2_ conditions across timepoints.

### Modulation of glucose and oxygen to endogenous-like cortical levels impact growth dynamics

To assess how *in vitro* culture conditions affect metabolic state of human cortical cell types, we adjusted the concentration of key nutrients, glucose and oxygen, to endogenous-like quantities. Standard glucose concentrations in culture media are 17.5 - 25mM; endogenous concentrations in amniotic fluid and fetal blood are approximately ⅕ the concentration ^34–37^. We steadily decreased glucose from day 9-18 to a final concentration of 5mM, which then remained consistent throughout culture duration. Additionally, while *in vitro* cultures are routinely maintained at atmospheric oxygen concentrations (20%), the oxygen tension in the brain, particularly during RG expansion periods, is significantly lower^38^. We modulated O_2_ concentrations to 8% to mimic physiologic levels (Figure 1A). We confirmed that 8% O_2_ did not induce expression of hypoxia genes (*HIF1a, BNIP3*), however a decrease in some glycolysis genes was observed (*PGK1, ENO1*) (S Figure 1D-E). These data support that oxygen and glucose collaborate to regulate metabolic state.

To explore the interaction between nutrient and O_2_ accessibility on neurodevelopmental regulation, we cultured forebrain organoids under standard (20%) and physiologic (8%) O_2_ and standard (18mM) and physiologic (5mM) glucose for 10 weeks (Figure 1A). Both glucose and O_2_ availability impacted organoid growth dynamics. Physiologic levels modestly increased the organoid growth rate during the first four weeks, however by six weeks, resulted in significantly decreased growth rate leading to smaller organoid size (Figure 1B, S Figure 1F). Given morphological observations, we evaluated if physiologic conditions have metabolic consequences by assessing extracellular flux across development. During routine media changes, the ‘spent’ media, including extracellularly secreted metabolites, were isolated and normalized to control media from the same condition. We observed that under 5mM glucose, there was a small reduction of glucose consumption during peak neurogenic periods (week 6-10), particularly compared to 8% O_2_ conditions (Figure 1C, S Figure 1G). We observed dynamic changes in lactate production; 18mM 8% O_2_ increased quantities of lactate, compared to 5mM glucose conditions at either O_2_ condition (Figure 1C, S Figure 1G). This data suggests a shift from increased anaerobic glycolysis, producing lactate, under 18mM conditions throughout neurogenesis, particularly at 8% O_2_. As higher glucose increased organoid size, yet lower O_2_ had higher glycolytic activity per cell (normalized to organoid size), these data indicate different metabolic programs are regulated by glucose vs O_2_ that impact cellular activities key for development. Abundance of some glycolysis enzymes were also impacted by changes in nutrient access, supporting the need for further functional interrogation (S Figure 1H-I).

### Untargeted metabolomics reveal distinct bioenergetic programs under physiologic conditions

To characterize metabolic activities across development, we analyzed intracellular metabolites from organoids grown under different conditions using untargeted metabolomics with a liquid chromatography mass spectrometry (LC-MS) system. Organoid samples were collected at weeks 6 and 10, given previous growth and metabolic observations (Figure 1B-C), and key progenitor and neuronal cell types expanded during these periods ^29^. We calculated metabolite z-scores to compare differences between 18mM vs 5mM glucose and 20% vs 8% O_2_ conditions (Figure 1D, S Figure 2A-B). Under standard 18mM glucose, we observed elevated metabolites corresponding to glycolysis (glucose, lactate), glycogen (glucose 1-phosphate, galactose), nucleotide sugars, and pentose phosphate (ribose phosphate, phosphoribosyl pyrophosphate) pathways related to the breakdown of glucose and fructose-6P (Figure 1D-E, S Table 1). In contrast, 5mM conditions displayed increased pyruvate metabolites, including TCA (citrate, malate) and related amino acids (alanine, isoleucine, arginine, aspartate, asparagine). Pathway analysis further confirmed these changes and highlighted differences in glucose-related versus amino acid pathways in 18mM versus 5mM conditions (Figure 1E, S Table 2). Differences in energetic pathways suggest primary utilization of glycolysis in 18mM, while 5mM conditions may utilize other catabolic programs, via mitochondria, to produce energy.

**Figure 2:**
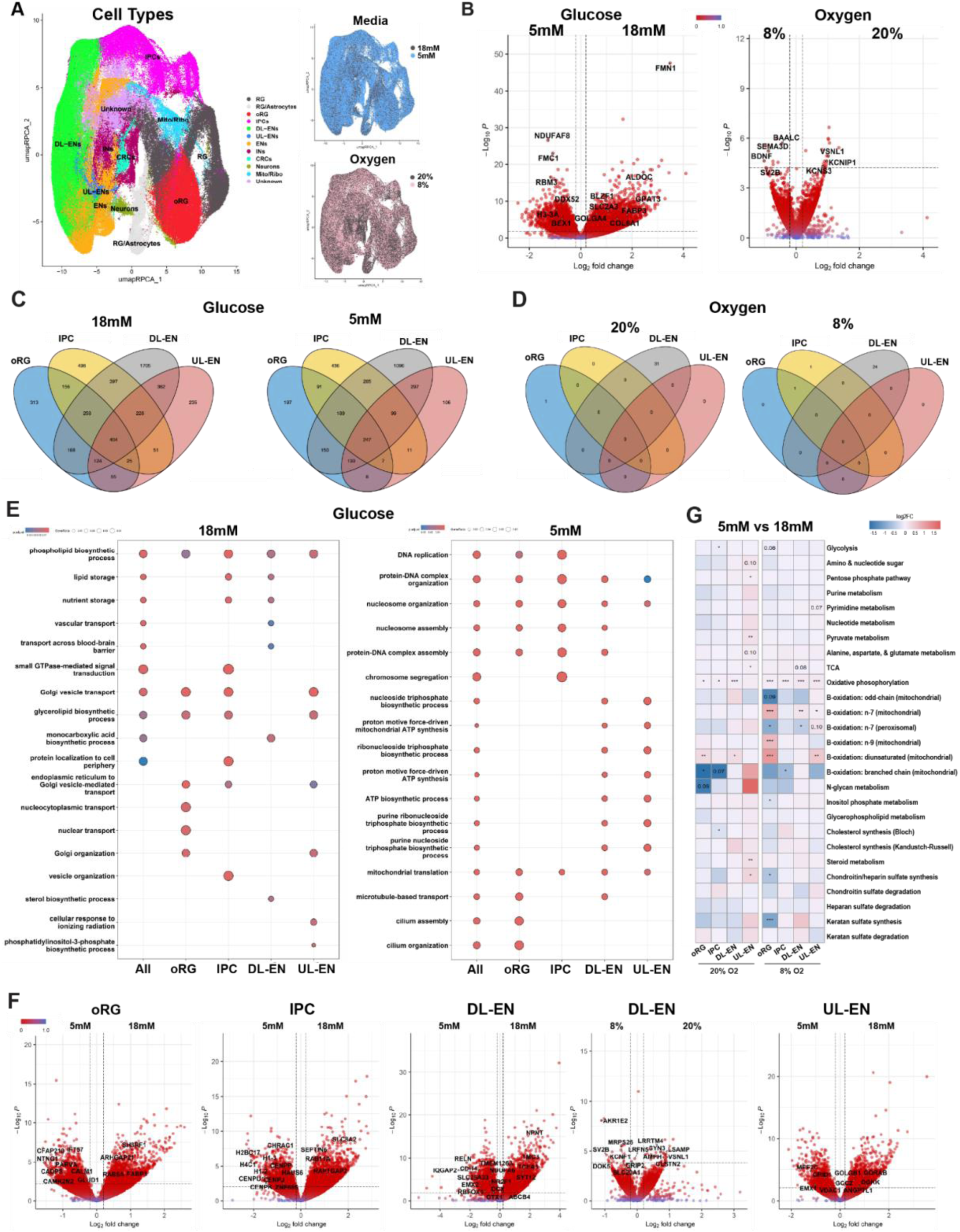
Sc-RNA-seq reveals molecular changes to cortical cell types as a consequence of physiological culture conditions. **A)** UMAP of organoid cells captured for sc-RNA-seq annotated by cell type, media, and O_2_ condition. B) Volcano plots of significant DEGs between 5mM vs 18mM glucose and 8% vs 20% O_2_ conditions across all cell types. C) There is both unique and shared overlap of DEGs for each glucose condition (18mM, 5mM) within cell types of interest: oRG, IPC, DL-EN, and UL-EN . D) There is no overlap of DEGs across cell types for O_2_ conditions (20%, 8%). Most DEGs are in the DL-EN cell type. E) Dotplots of GOTerm Enrichment from DEGs across all cell types of interest (oRG, IPC, DL-EN and UL-EN) combined (leftmost) or within each respective cell type (per 18mM or 5mM glucose condition). Shared programs are primarily metabolic regulators, while unique programs indicate cell-specific changes. F) Volcano plots of 5 vs 18mM glucose conditions in oRG, IPC, DL-EN, and UL-EN and 8 vs 20% O_2_ demonstrate cell type-specific molecular changes. G)METAFlux analysis demonstrates differences between 5mM glucose conditions compared to 18mM control, across O_2_ conditions, within a respective cell type.

O_2_-driven metabolic changes were broader, with organoids cultured under standard 20% O_2_ exhibiting an overall increase in metabolic activity. Citrate, isocitrate, malate, and related mitochondrial amino acids (threonine, asparagine, glutamine) were increased compared to 8% conditions (Figure 1D, S Figure 2A, C). These data suggest that while we intended to mimic tissue-level quantities of O_2_, perhaps the quantity was insufficient for O_2_-intensive programs, as occur in mitochondria, despite not triggering hypoxia (S Figure 1E). However, changes did not increase cytosolic bioenergetic programs, such as, glycogen, glycolysis, or pentose phosphate (Figure 1D, F, S Table 1-2). Taken together, this global pathway suppression indicates less metabolically active cells in the 8% conditions, which may underlie previously observed differences in organoid growth. These phenotypes were more pronounced at week 10, compared to week 6, suggesting temporal shifts in O_2_ utilization that correspond with neuronal differentiation. In general, while we observed subtle shifts in specific metabolites between weeks 6 and week 10 (Figure 1E-F, S Figure 2B-C), most pathway changes observed in glucose and O_2_ conditions were consistent across time points.

### Molecular phenotyping of organoids under physiologic glucose and oxygen

To assess molecular consequences of metabolic alterations on neurodevelopment, we isolated organoids from weeks 6 and 10 and captured live cells for single cell RNA sequencing (sc-RNA-seq). After quality control filters (S Table 3), 248,661 cells were analyzed from organoids derived from three PSC lines (H1, 13234, H21792) that were cultured under 20% or 8% O_2_ and 18mM or 5mM glucose. Sequenced samples were aligned to the reference genome, quality control performed, and samples integrated prior to clustering. Clustered samples were annotated^29,33^ and granular annotations were then combined for clarity of cell type classification (S Figure 3A). Based on gene expression (S Table 4), our final clustered dataset was divided into progenitor subtypes: RG, oRG, IPCs; and neuronal subtypes: deep layer (DL-) excitatory neurons (EN), upper layer (UL-) EN, pan EN, inhibitory interneurons (IN), and Cajal-Retzius cells (CRCs), as well as one cluster driven by mitochondrial/ribosomal genes (Figure 2A, S Figure A-E). Cells across media, oxygen conditions, cell line, and time point were observed across clusters (Figure 2A, S Figure 3A). To ensure that conditions did not have unintended impact on cortical differentiation efficiency, we verified anticipated marker expression for cortical progenitors (*PAX6, FOXG1*), division (*KI67*), and specific subtypes, including neurogenic IPC (*EOMES*) and human-expanded oRG cells (*HOPX*) (S Figure 3B-E). We also observed anticipated differentiation of neurons (*NEUROD6*), into subtypes of early-born DL-EN (*BCL11B*), later-born UL-EN (*SATB2*), and inhibitory INs (*DLX5*) (S Figure 3D-E). To confirm that glucose was the key substrate driving metabolic differences, sc-RNA-seq analysis included assessment of different basal media composition containing 5mM glucose (DMEM/F12, RPMI, Plasmax; S Figure 3F-G). We observed more differentially expressed genes (DEGs) shared across media types than were unique to each medium. Gene Ontology (GO) was performed on overlapping and unique genes per media, which indicated shared programs when all 5mM media were evaluated together (Figure 2E, S Figure 3F-G). Due to phenotypic similarities, we conducted subsequent analysis of the media conditions by pooling DMEM/F12, Plasmax, and RPMI media into one 5mM glucose group.

**Figure 3:**
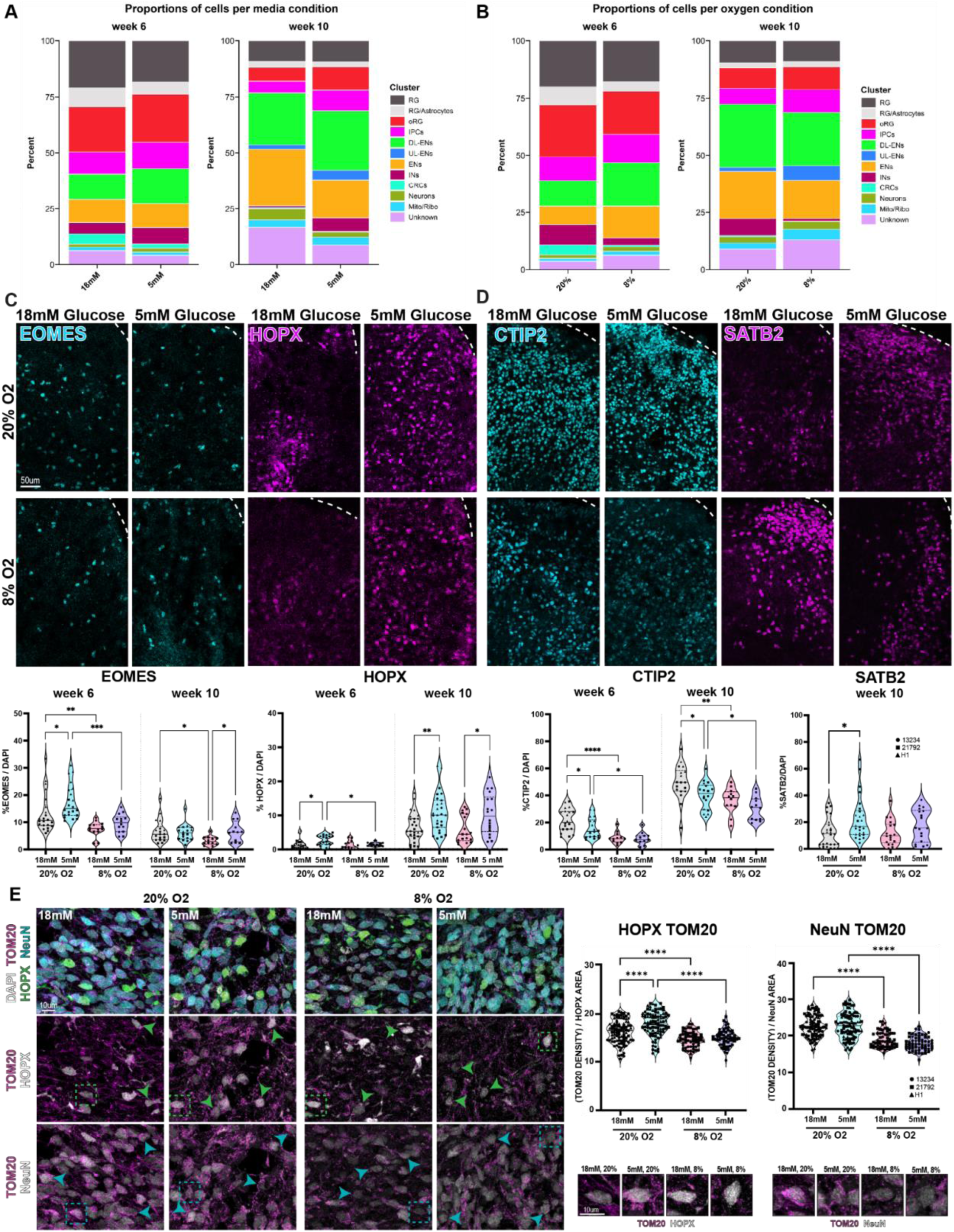
Physiological glucose and oxygen regulate fate via mitochondrial metabolism. **A)** Cell type proportions captured per glucose/media condition from week 6 and week 10 organoids (n=2 organoids each from n=3 iPSC lines/condition/timepoint). **B)** Cell type proportions captured per O_2_ condition from week 6 and week 10 organoids. **C)** EOMES+ IPC decrease in 8% O_2_ (Ordinary one-way ANOVA: Week 6: 18mM 20% vs 5mM 20%: *p<0.0322; 18mM 20% vs 18mM 8%: **p<0.0097; 5mM 20% vs 5mM 8%: ***p<0.0004; Week 10: 18mM 20% vs 18mM 8%: *p<0.0261; 18mM 8% vs 5mM 8%: *p<0.0443). HOPX+ oRG increase in 5mM glucose conditions (Ordinary one-way ANOVA: Week 6: 18mM 20% vs 5mM 20%: *p<0.0170; 5mM 20% vs 5mM 8%: *p<0.0232; Week 10: 18mM 20% vs 5mM 20%: **p<0.0090 ; 18mM 8% vs 5mM 8%: *p<0.0282). Points represent images from n=4 organoids per differentiation batch from n=3 iPSC lines/condition/timepoint. **D)** CTIP2+ DL-ENs decrease in 8% O_2_ (Ordinary one-way ANOVA: Week6: 18mM 20% vs 5mM 20%: *p<0.0253; 18mM 20% vs 18mM 8%: ****p<0.0001; 5mM 20% vs 5mM 8%: *p<0.0179; Week 10: 18mM 20% vs 5mM 20%: *p<0.0287; 18mM 20% vs 18mM 8%: **p<0.0053; 5mM 20% vs 5mM 8%: *p<0.0344). SATB2+ UL-ENs increase in 5mM glucose (Ordinary one-way ANOVA: Week 10: 18mM 20% vs 5mM 20%: *p<0.0160). **E)** TOM20 integrated density in HOPX+ oRGs decreases in 8% O_2_, but increases in 5mM glucose under 20% O_2_ (Ordinary one-way ANOVA: 18mM 20% vs 5mM 20%: ****p<0.0001; 18mM 20% vs 18mM 8%: ****p<0.0001; 5mM 20% vs 5mM 8%: ****p<0.0001). TOM20 integrated density in NEUN+ neurons decreases in 8% O_2_ but is unaffected by glucose (Ordinary one-way ANOVA: 18mM 20% vs 18mM 8%: ****p<0.0001; 5mM 20% vs 5mM 8%: ****p<0.0001). Points represent individual cells from n=4 organoids from n=3 iPSC lines/condition.

### Environmental conditions impact global metabolic programs

We evaluated how nutrient access impacts the molecular state of organoid cells by assessing DEGs across conditions. Given the more significant changes at week 10 in the LC-MS dataset (Figure 1D-E), and increase in DEGs in the week 10 samples (6,562) compared to week 6 (628) (S Figure 3G), we primarily focused our evaluation on week 10 organoids. Glucose concentration drove global metabolic changes across organoid cell types, which had a greater impact on gene expression than O_2_ (Figure 2B, S Table 5). Across all cell types, under 18mM glucose conditions, there was consistent increase in genes regulating glucose uptake/breakdown (*ALDOC, ARRDC4, SLC2A3, C1QTNF12*), glycerophospholipid/phospholipid biosynthesis (*DGKG/E, LPCAT4, GPAT3, ENPP2, FABP3, FIG4, FITM1, GDPD3*), Golgi regulation (*BLZF1, HACE1, GOLGA3/4*), and morphology-regulating extracellular matrix (ECM)/cytoskeleton (F*MN1, COL6A1/2, TUBA4A, ADAMTS2, MATN*1). Whereas under 5mM glucose conditions, there was higher expression of genes related to mitochondrial respiration (*FMC1, NDUFAF8/11, MRPL20, SELENOH, TOMM22, COX20, NDUFB2, SLC25A33, GDAP1, GATD3)*, chromatin accessibility (M*LLT1, RCBTB2, HMGN1/2, BEX1,PCLAF, H3-3*A), and RNA binding (*RBM17, GPATCH4, RBM3, HNRNPA3, SYNCRIP, HNRNPDL, DDX52*) across cell types (Figure 2B). The significance and level of expression differences between groups was lower under O_2_ conditions. Under 20% O_2_, DEGs indicated physiological function via calcium sensing/potassium channels (V*SNL1, KCNIP1, KCNS3*), while 8% O_2_ conditions had increased expression of genes associated with neuronal growth and synaptogenesis (*SV2B, BAALC, SEMA3D, BAALC, BDNF*). Changes in glucose concentration appeared to broadly influence cellular metabolism, morphology, protein processing, and gene expression, while changes in oxygen concentration influenced regulators of neuron development and function.

To disentangle the cell-specific impact of nutrient perturbations, we assessed DEGs in cortical cell types of particular interest, oRG, IPC, DL-EN and UL-EN, and compared overlapping expression to decipher cell-specific features (Figure 2C-D). For glucose conditions, 18mM shared 404 genes and 5mM shared 247 genes; we then performed gene ontology (GO) on shared DEGs across cell types and observed global impact on metabolic state. Under 18mM glucose, all cells had increased Golgi-regulatory and glycophospholipid biosynthesis programs, suggesting impact on protein packaging, glycosylation, and transport. In contrast, 5mM glucose drove mitochondrial metabolism and ATP synthesis indicating transition to OXPHOS (Figure 2E, S Table 6-7), as anticipated from metabolomics observations (Figure 1D). For O_2_ conditions, we observed no overlap of DEGs across cell types, suggesting that while glucose availability has global consequences across cell types, oxygen had a modest impact that was specific to DL-ENs (Figure 2C-D).

### Cell-specific consequences of metabolic shifts

We sought to explore the impact of metabolic shifts on discrete developing cell types. Given the importance of oRG cells for human cortical expansion^39–42^, we evaluated how glucose impacted their molecular state (Figure 2E-F, S Table 8). Under standard 18mM glucose, we observed increased expression of genes associated with Golgi vesicle transport (*RAB1A/5A, SH3RF1, ARHGAP2*1), as well as global metabolic programs described above. Under physiologic 5mM glucose, oRG cells expressed genes associated with cytoskeletal movement/transport (*PARVA, NTNG1, MYL12B, MFSD4A*), cilium assembly (*IFT57, CFAP210* ), and receptor/channel activity (*CALM1, CADPS, GLUD1, CAMK2N2, SCN8A, CAMK2N2*) (Figure 2F). These data suggest that oRGs are sensitive to glucose concentration, which may impact signaling, nutrient sensing, and migratory capacity ^22^.

Neurogenic IPCs are essential transit amplifying cells that promote neuronal expansion. We observed that 18mM glucose supported increased expression of GTPase/CDC42 signaling (*RAP1GAP2, SEPTIN6, RAB3A, RAB11A*), calcium-mediated signaling (*SLC8A2, CLU*), and Golgi vesicle transport (R*AB6B, RAB1A, GOLGA3, GOLGA5*), suggesting shifts in morphology and signaling. Whereas, under 5mM glucose, IPCs had increased expression of genes associated with chromatin accessibility (*CHRAC1, H2BC17, H4C1, H1-3, H1-2, ANP32B, ZNF695, ACTL6A*) and cell division (*MTUS1, HAUS6, CENPP, CENPK, CENPJ, CENPU*) (Figure 2E-F, S Table 9) suggesting increased self-renewal.

The neuronal progeny of IPCs, DL-EN, have increased expression of genes regulating ionic and carboxylic acid transport (T*MC3, ABCB4, ADORA2A-AS1, ANXA11, CAMK2G, TCEA3, SYT12*) and ECM (A*BI3BP, P4HA2, NPNT*) under 18mM that suggest increased signaling and connectivity. Under 5mM glucose, in addition to increased expression of cell-specific mitochondrial genes (*SLC25A33, NDUFA6, TMEM126A, SLC25A39, ATP6V1G1, NDUFB9*), DL-EN expressed markers associated with neuron projection and migration (*RBFOX1, RELN, CDH4, IQGAP2, DCX, NRG1, CUX2, PLXNB1*) (Figure 2E-F). Interestingly, the DL-EN was the only population with phenotypes associated with O_2_ availability. DL-EN cultured under 20% O_2_ expressed genes regulating synapse formation, neuron morphology (S*YN3, LRRTM4, VSNL1, AMPH, LSAMP, CLSTN2*), and ionic signaling/activity (CPNE5, KCNIP1, GRIK5), whereas 8% O_2_ were associated with catabolic processes (*AKR1E2, MRPS26*) and vesicle exocytosis/neuron transmission (S*V2B, GRIP2, DOK5, SLC20A1, KCNF1*) (Figure 2D, F; S Table 10). These data suggest DL-ENs may be uniquely sensitive to oxygen and glucose availability during their differentiation.

The human-expanded progeny of oRG, the UL-ENs, had modest cell-specific changes associated with glucose concentration. Under 18mM glucose, DEGs are associated with phospholipid/glycerolipid metabolic programs (*ABHD18, ABHD5, ANGPTL1, DGKK*) and Golgi processing of proteins (*GORAB, GOLGB1, GCC2*). In 5mM glucose, UL-ENs had increased expression of respiration (VDAC1, NDUFB9) and neuronal maturation-associated genes (*MEF2C, GRID1*) (Figure 2E-F, S Table 11). Interestingly, under 5mM conditions, DL-ENs (*OTX1, NR2F1, EMX2*) and UL EN (*EMX1*) expressed markers of cortical areal commitment, suggesting mitochondrial metabolism may also play a role in patterning programs ^29^.

### Metabolic evaluation of cell-specific changes using transcriptomics

To define more nuanced differences between metabolic states in discrete cell types across conditions, we employed METAflux ^43^ analysis (Figure 2G, S Table 12). A random subset of cells per condition, across all PSC lines, was used to define flux score per metabolic pathway. The 5mM glucose condition was compared to standard 18mM glucose, within each respective O_2_ condition. Mitochondrial OXPHOS had a small, but significant increase, across cell types in 5mM conditions, compared to 18mM conditions, regardless of O_2_ concentration. Supportively, there were also modest increases in TCA and pyruvate metabolism in UL-ENs and decreased glycolysis in IPCs under 5mM conditions.

Dynamic regulation of beta oxidation of fatty acids was observed across cell types and conditions, with most alterations in the progenitors. Under 5mM glucose, oRG had increased flux of mitochondrial fatty acid oxidation (ATP production), but decreased peroxisomal fatty acid oxidation (lipid metabolism), which was most pronounced under 8% O_2_ ^44^. Recent studies have revealed the unique importance of mitochondrial beta-oxidation in temporal pacing of human-specific neurodevelopment ^45^. We also observed N-glycan metabolism trending down in progenitors but up in ENs; as well as increasing trends of alanine, aspartate, and glutamate metabolism, amino acid and nucleotide sugars, and pyrimidine metabolism specifically in the UL-ENs. These pathways align with metabolomics phenotypes observed in week 10 organoids, suggesting that their increased utilization may drive UL-EN expansion. These results corroborate functional LC-MS observations and DEGs using alternative computational approaches to define cell-specific programs regulated by nutrient availability (Figures 1D, 2E). Collectively, these data support that nutrient access regulates the global metabolic state of organoids, but subtle shifts in cell-specific activities can be evaluated using estimations of flux. Data from both bulk and cell-specific analyses consistently show that glucose concentration regulates metabolic pathway usage and O_2_ availability amplifies those changes.

### Oxygen and glucose impact cell fate decisions concurrent with mitochondrial dynamic shifts

To assess the longitudinal impact of global metabolic shifts on fate decisions, we evaluated cell composition in organoids cultured under physiologic nutrient conditions from sc-RNA-seq data. We observed a proportional increase in captured oRG cells under 5mM glucose (red cluster) and their progeny, UL-EN (blue cluster) (Figure 3A), suggesting support for cortical cell subtypes expanded in the human brain. In contrast, physiological levels of O_2_ (8%) did not impact major cell type compositional changes; subtle shifts in different EN clusters and mitochondrially-defined clusters were observed (Figure 3B).

To quantify hypothesized fate shifts observed in RNA capture, we used immunohistochemistry (IHC) to assess progenitor populations, EOMES+ IPCs and HOPX+ oRG cells. At week 6, physiologic 5mM glucose resulted in a small increase in EOMES+ IPCs, whereas physiological 8% O_2_ significantly reduced numbers (Figure 3C). By week 10, 8% O_2_ continued to decrease IPC production with 5mM glucose modulating impact. In contrast, 5mM glucose promoted production of HOPX+ oRG cells across timepoints. 8% O_2_ impaired this production at week 6, but had no impact at week 10 (Figure 3C).

To determine the consequences of progenitor pool diversification, we evaluated organoids for the early-born CTIP2+ DL-EN and later-born SATB2+ UL-EN. Since physiologic 8% O_2_ impairs neurogenic IPC production, we assessed the impact on DL-EN progeny and observed a corresponding reduction across physiologic glucose and O_2_ conditions (Figure 3D). Given the expansion of oRG cells under 5mM glucose, we sought to assess the production of their SATB2+ UL-EN progeny. Supporting sc-RNA-seq data, we observed increased production of SATB2+ neurons under physiologic glucose (Figure 3D).

To validate whether cell composition differences were a consequence of proliferation changes, we quantified the presence of pHH3, a mitotic marker, and the broad RG marker, PAX6. Despite organoid growth rate differences (Figure 1C), we observed no change in pHH3+ mitotic cells and only a small reduction at week 6 in the number of PAX6+ RG when cultured at 5mM glucose and 8% O_2_ (S Figure 3H), suggesting minimal impact on proliferation. To verify alterations in size/composition were not due to cell death, we performed a TUNEL assay and observed no significant differences across culture conditions (S Figure 3I).

As 5mM glucose and 20% O_2_ promote the generation of oRGs and neurons, we wanted to evaluate whether phenotypes were driven by mitochondrial activities observed in our previous analyses (Figure 1D, 2E). We performed IHC of TOM20 to examine mitochondrial content in HOPX+ oRGs and NEUN+ neurons at week 10 (Figure 3E). Mitochondria were O_2_-sensitive in both populations, indicating both oRGs and EN utilize aerobic programs. In HOPX+ oRGs, TOM20 intensity was significantly higher in 20% O_2_ and, importantly, increased in 5mM glucose conditions, suggesting that decreased glucose availability may activate mitochondrial programs affiliated with oRG development. In NEUN+ neurons, TOM20 signal intensity also significantly decreased under both 8% O_2_ conditions, but was unaffected by glucose concentration indicating decreased reliance on glucose-derived energy. IHC validation studies suggest that mitochondria programs are suppressed in 8% O_2_, despite predictions of higher OXPHOS flux in both O_2_ conditions in our METAflux analysis (Figure 2G). These discrepancies may suggest lower mitochondria content in 8% O_2_ triggers compensatory OXPHOS flux in remaining mitochondria to meet energy demands, potentially at the expense of biosynthesis. Decrease in biosynthesis pathways aligns with reduced organoid growth in 8% O_2_. Thus, appropriate nutrient conditions are essential to support the metabolic requirements of cortical populations as they transition to oxidative phenotypes.

### Metabolic transition from anaerobic to aerobic glycolysis drive neurodevelopmental changes

To functionally evaluate the impact of glucose utilization on identified fate changes, we inhibited glycolysis enzymes in week 6 organoids, during progenitor diversification (Figure 4A). ShRNAs were designed against TPI1 and LDHA and introduced at week 6 using an established electroporation paradigm ^46^. Organoids were evaluated 1-week later after robust fluorescent reporter expression; knockdown efficacy was confirmed and there was no impact on viability (S Figure 4A-C). We assessed the cell autonomous impact on identity in GFP+ cells and observed that TPI1 knockdown trended toward an increase in CTIP2+ DL-EN, compared to control shRNA (Figure 4B). In contrast, LDHA knockdown specifically promoted increase in HOPX+ oRG (Figure 4B, S Figure 4D).

**Figure 4:**
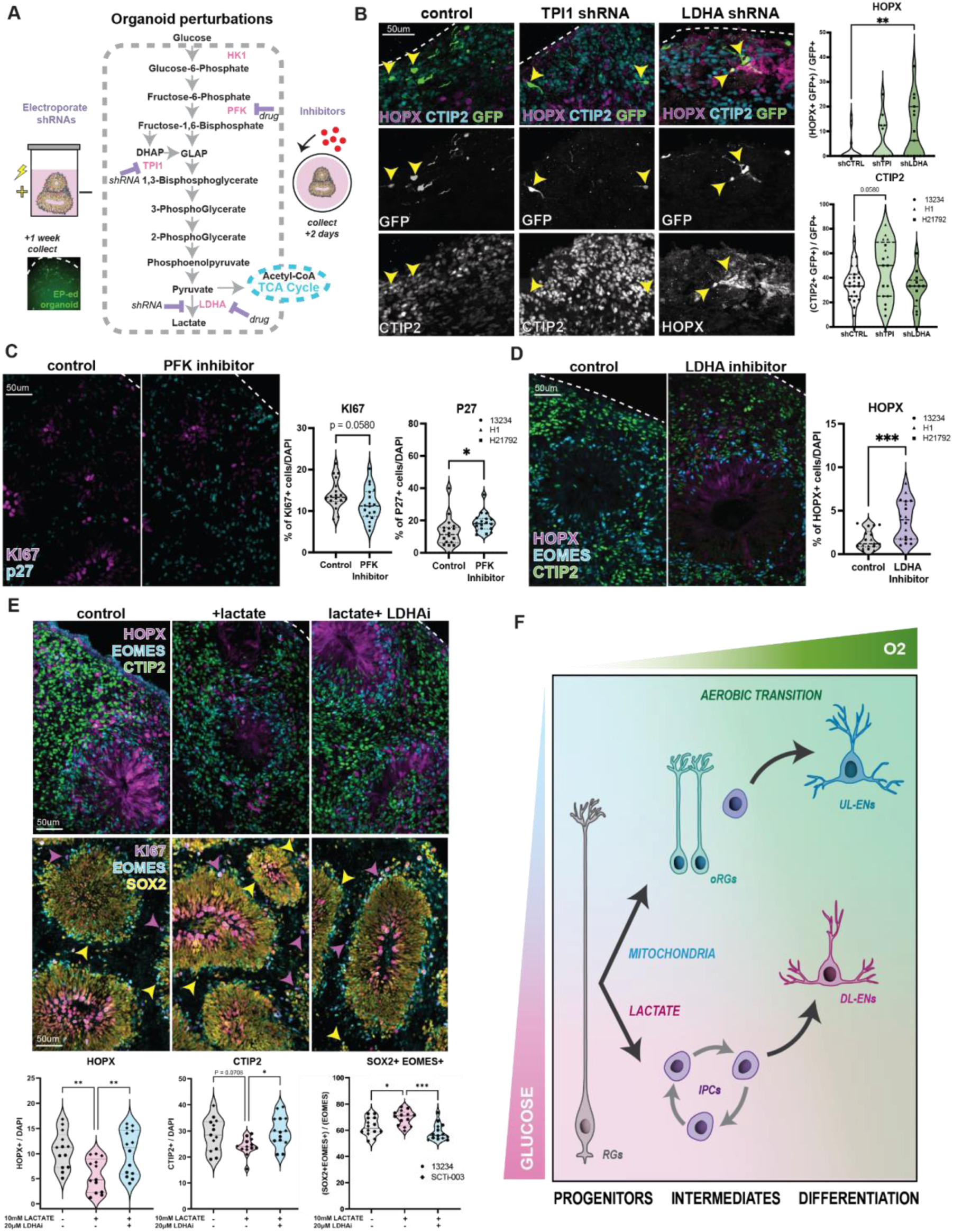
Total glycolysis inhibition promotes neurogenesis, while anaerobic glycolysis/lactate inhibition promotes oRG production. **A)** Glycolysis schematic with metabolites (black), enzymes (pink) and perturbations in this study (purple). Organoids are differentiated for 6 weeks before exposure to pharmacological perturbations for 2 days or shRNA electroporation and collected 7 days later. **B)** Week 6 organoids electroporated with shTPI1 or shLDHA altered cell fate. GFP+ HOPX+ oRG cells increase after LDHA shRNA electroporation compared to control shRNA (One-way ANOVA, **p<0.0048). GFP+ CTIP2+ DL-ENs increase after TPI1 shRNA electroporation compared to controls (One-way ANOVA, **p<0.0048). Points represent images from n=3 organoids per differentiation batch from n=3 iPSC lines/condition. **C)** Week 6 organoids treated with PFK15 have increased P27+ post-mitotic cells (Unpaired student’s t-test: *p<0.0229) and a trend toward a decreased KI67+ cycling cells (unpaired t test: p=0.058). Points represent n=3 organoids per differentiation batch each from n=3 iPSC lines/condition. **D)** Week 6 organoids treated with GSK2837808A have increased HOPX+ oRGs (Shapiro-Wilk normality: p<0.0226; Mann-Whitney test, ***p<0.0007). Points represent n=3 organoids per differentiation batch from n=3 iPSC lines/condition. **E)** Week 6 organoids treated with 10mM lactate have decreased HOPX+ oRGs that are rescued with 20uM GSK2837808A (one-way ANOVA, control vs lactate: **p<0.0059; lactate vs lactate+inhibitor: **p<0.0074). Lactate yielded a trending decrease in CTIP2+ DL-ENs that normalized in combination with GSK2837808A (one-way ANOVA, control vs lactate: p=0.0708; lactate vs lactate+inhibitor: p<0.0181). Lactate increased SOX2+EOMES+ IPCs that normalized with GSK2837808A (one-way ANOVA, control vs lactate: *p = 0.0123; lactate vs lactate+inhibitor: ***p = 0.0001; control vs lactate+LDHAi: p=0.0871). Points represent n=3 organoids per differentiation batch from n=2 iPSC lines/condition. **F)** Schematic summarizing the effects of glucose and oxygen on human neural cell types.

To corroborate findings, we treated week 6 organoids with pharmacological inhibitors of PFK, to decrease glycolysis pathway function, or LDHA, to specifically inhibit conversion of pyruvate to lactate, as occurs in anaerobic glycolysis. Organoids were treated with inhibitors for 2 days, before collecting for analysis (Figure 4A, S Figure 4E). As anticipated from previous studies ^8,23^, total inhibition of glycolysis through PFK, led to increased p27+ post-mitotic differentiated cells and a trend toward reduction of KI67+ proliferating cells (Figure 4C). Inhibiting glycolysis impacted differentiation rate, resulted in a small increase in TUNEL+ cell death, but did not affect the production of HOPX+ oRG, EOMES+ IPCs, or CTIP2+ DL-ENs (S Figure 4F). In contrast, treatment with an LDHA inhibitor resulted in a specific increase in HOPX+ oRG cells, with no impact on the fate of other neural cell types or proliferation/differentiation programs (Figure 4D, S Figure 4G).

Data suggests that total suppression of glycolysis promotes neuronal differentiation, while targeted inhibition of LDHA, and thus lactate, is required for oRG production. This observation could be a consequence of anaerobic to aerobic glycolysis switch within oRG or because of environmental lactate concentration changes, due to metabolic switch in neighboring cell types. To determine lactate sufficiency to impact oRG development, we added 10mM lactate at week 6 for 2 days and quantified changes in cell fate. Increased lactate concentrations decreased the proportion of HOPX+ oRGs, in addition to a trending decrease in CTIP2+ DL-ENs (Figure 4E). LDHA inhibition, in combination with lactate addition, rescued these phenotypes. EOMES+ IPCs and other neuronal populations were unaffected at week 6 (S Figure 4H-I). We then assessed whether lactate promoted immature phenotypes. While no change in KI67+ proliferating cells or PAX6+ RG were observed, the proportion of proliferating SOX2+EOMES+ and KI67+EOMES+ IPCs were significantly increased (Figure 4E, S Figure 4J). These data suggests lactate promotes immature IPC states; inhibition promotes oRG specification. Together, study data indicate that appropriate access to environmental nutrients impacts the metabolic state regulating the fate of neural cells during human neurogenesis (Figure 4F).

## Discussion

Our studies evaluated how changes in nutrient accessibility in the cortical environment impact developmental programs. Previous studies demonstrate that glycolysis drives RG proliferation and transition to cellular respiration for neuronal differentiation ^8,23,25^. However, the impact of metabolism on developmental state in the resolution of expanded progenitor and neuronal subtypes in the human brain has not been extensively explored. While physiologic changes in glucose and O_2_ availability affect metabolic activity, we do not observe changes in proliferation dynamics. However, under glycolysis suppression, via inhibitors of PFK or TPI1, differentiation is promoted, as expected. We observed that lower O_2_ levels decreased organoid growth rates, metabolic activity, and suppressed neuronal differentiation. Size differences may be due to decreased neuronal production, a known phenotype in microcephaly ^47^. Oxygen-related metabolic changes predominantly affected amino acids (glycine/serine/threonine), which have demonstrated roles for neurotransmitter synthesis and mitochondrial programs that support neuron functions^48^. Changes in neuronal mitochondria content and RNA programs in response to O_2_ concentration is indicative of their oxidative phenotype to meet high energy demands. To balance oxidative stress from high mitochondria metabolism, neurons utilize glucose through the PPP, to generate reducing agents ^49,50^. The increase in DL-ENs, following TPI1 inhibition, may also be due to the diversion of glucose into the oxidative branch of the PPP, as suggested by recent metabolic atlasing studies of primary human cortex during late neurogenic/early gliogenic periods ^51^. Together, results suggest that O_2_ regulates metabolic phenotypes that are essential for neuronal production in cortical development.

While a switch from anaerobic glycolysis to aerobic respiration occurs during neural stem cell differentiation into post-mitotic neurons, the intermediate pathways that govern progenitor transitions are less understood. Our results suggest that glucose availability regulates glycolytic versus mitochondrial programs implicated in the diversification of discrete cell populations. While direct branches of glycolysis were elevated in standard 18mM conditions, 5mM glucose-treated organoids showed increased anaplerosis and TCA functions. These activities in physiologic glucose closely resembled those of the 20% O_2_ conditions, suggesting that decreased glycolytic flux activates mitochondrial programs to produce ATP. The implications of increased aerobic capacity for regulating cell type specification are demonstrated by the concurrent increase of oRGs and UL-ENs in 5mM conditions. During their peak production, oRGs, under 5mM glucose, exhibited increased mitochondrial content and higher expression of genes involved in amino acid metabolism, catabolism, and mitochondrial function. This aligns with previous studies demonstrating that human-specific variants of oRG genes, particularly ARHGAP11B, contribute to their expansion by enhancing metabolism ^52,53^. The subsequent transition to primarily aerobic UL-ENs was demonstrated by similar increases in OXPHOS, mitochondrial organization, and redox homeostasis programs under 5mM conditions. Collective changes indicate that appropriate glucose levels enable more refined metabolic states that impact fate determination programs.

Transient changes in glucose reliance during neuronal lineage progression were further demonstrated by cell type-specific effects of different glycolysis inhibitors. While upstream suppression of glycolysis altered progenitor/proliferative versus post-mitotic states, the fate of glycolytic end-products exhibited differential effects on discrete progenitor populations. Specifically, the negative relationship between lactate and oRG production was demonstrated by LDHA knockdown, inhibition, and extracellular lactate supplementation. While increased mechanistic resolution of lactate regulation of oRG production should be explored in future studies, various works have demonstrated the impact of lactate on cellular morphology and migration via interactions with cAMP, FGF, and STAT3,^54–56^ signaling, which influence oRG development ^33^. Given the morphological programs upregulated in 5mM oRGs, which are key for their functions^57^, it is possible that higher lactate concentrations promote more immature states, preventing specialized subtype activities. This is supported by the lactate-driven increase in immature, proliferative IPCs during early neurogenesis and trending decrease in neuronal differentiation after short-term exposure. Lactate-driven effects could underlie the increased DL-ENs in 18mM conditions, potentially at the expense of UL-EN populations. Previous works have also demonstrated the role of lactate in early neurodevelopment by promoting self-renewal states via mitochondria fusion and triggering angiogenesis^8^. However, the specific effects of lactate on discrete oRG and IPC behaviors require further investigation to understand its regulatory role in human corticogenesis. Collectively, our findings demonstrate that glucose utilization regulates progenitor transitions that govern cortical laminae-specific expansion, while oxygen-dependent programs in the mitochondria drive neuronal differentiation.

Given the protracted duration of progenitor expansion leading to increased cell type heterogeneity in the human brain and the essential role of metabolism in neural tissue formation, these studies serve as an important step toward defining metabolic governance of temporal fate transitions. Observations highlight metabolic differences in the regulation of cell types expanded in the human brain, which may have increased vulnerability to changes in metabolism during gestation. Lack of glucose and O_2_ availability, either due to injury or genetic mutation, are associated with a range of neurodevelopmental disorders ^11,58–63^. Future studies can leverage identified changes to target metabolically-regulated disruption in patient iPSC lines to study the mechanistic origin of these conditions. Moving forward, a focus on functional assessments of real-time, cell-specific changes are needed to interrogate temporal transitions of developing cell types to gain new insights into the role of metabolism in disease.

### Limitations of this study

A limitation to this study is the lack of vasculature present within *in vitro* organoids that do not provide local sources of O_2_ and glucose. However, simplified iPSC models equip assessment of requirement and sufficiency of metabolic programs for developmental changes. Stem cell models also lack additional aspects of *in vivo* physiology and circuitry, despite their increased pliability to manipulate and quantify molecular programs.

## STAR Methods

### Key Resources Table

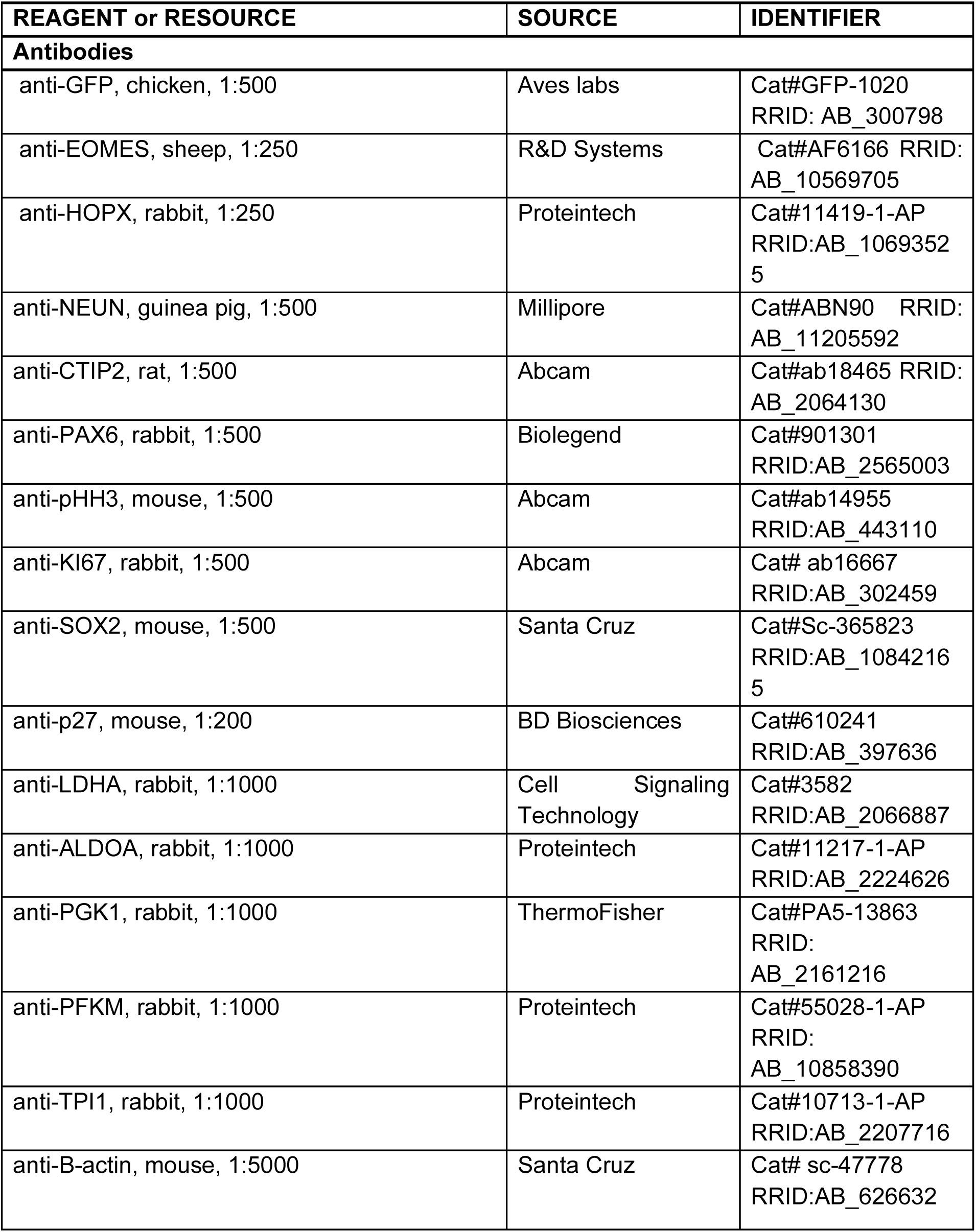
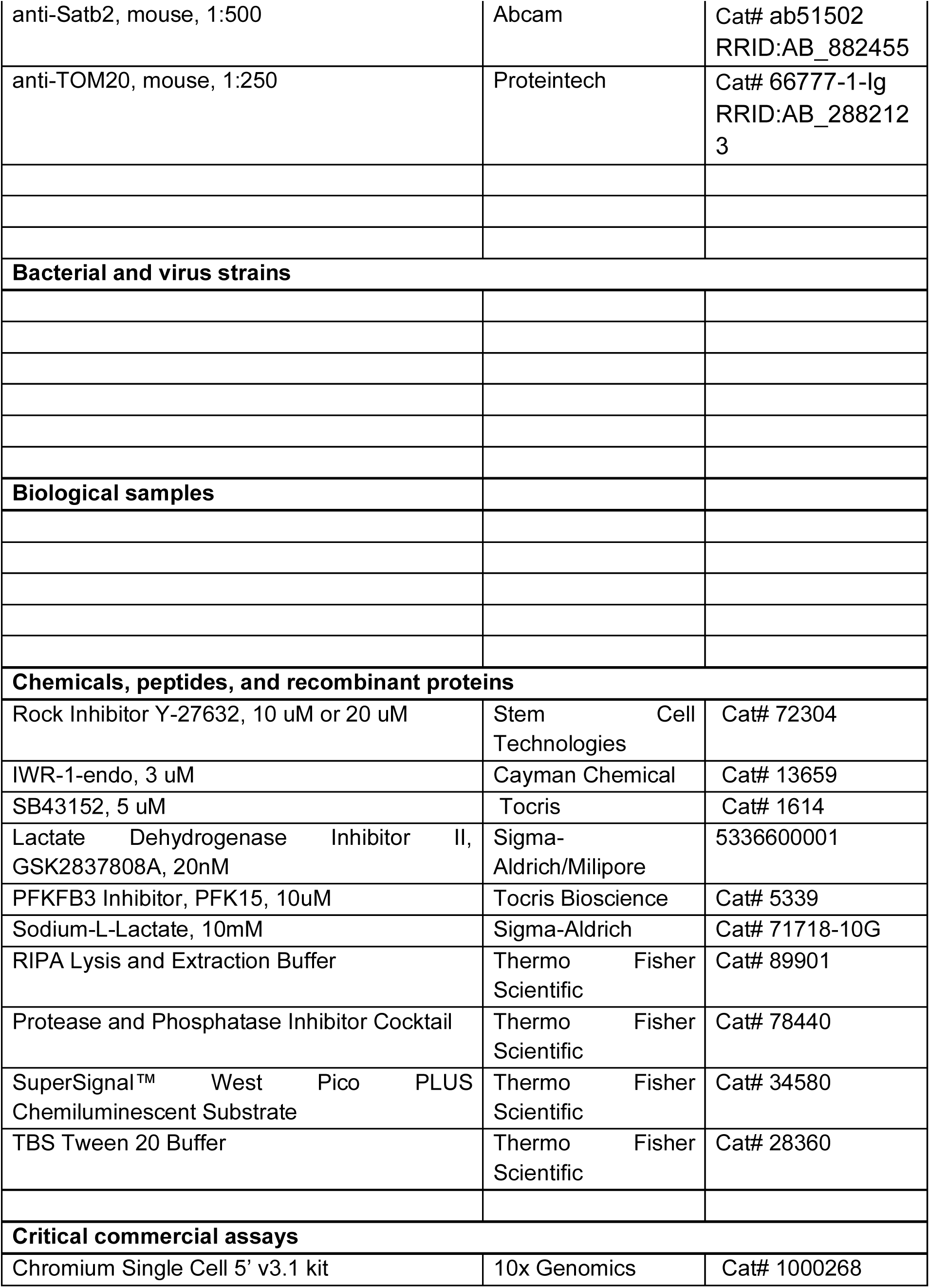
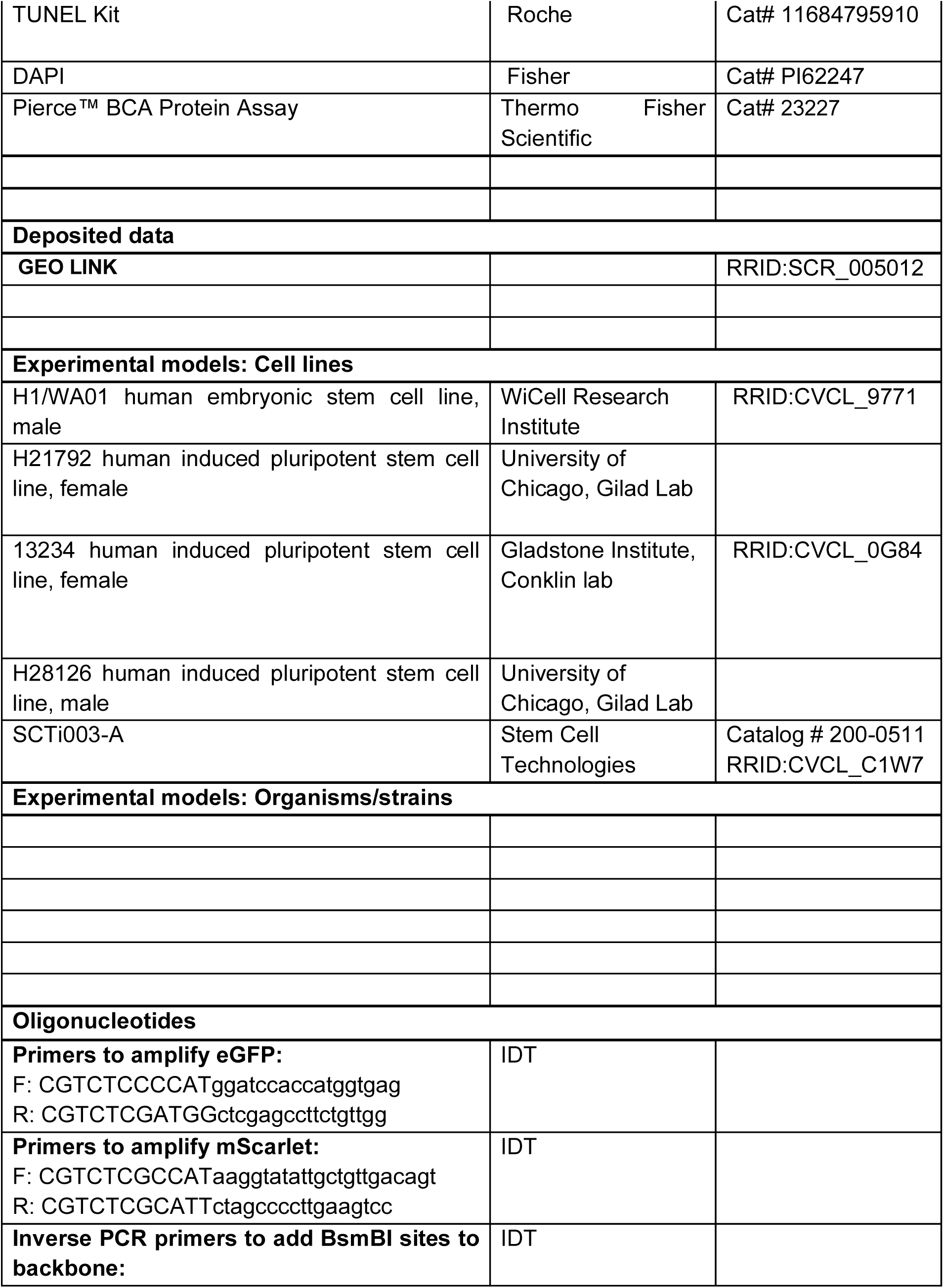
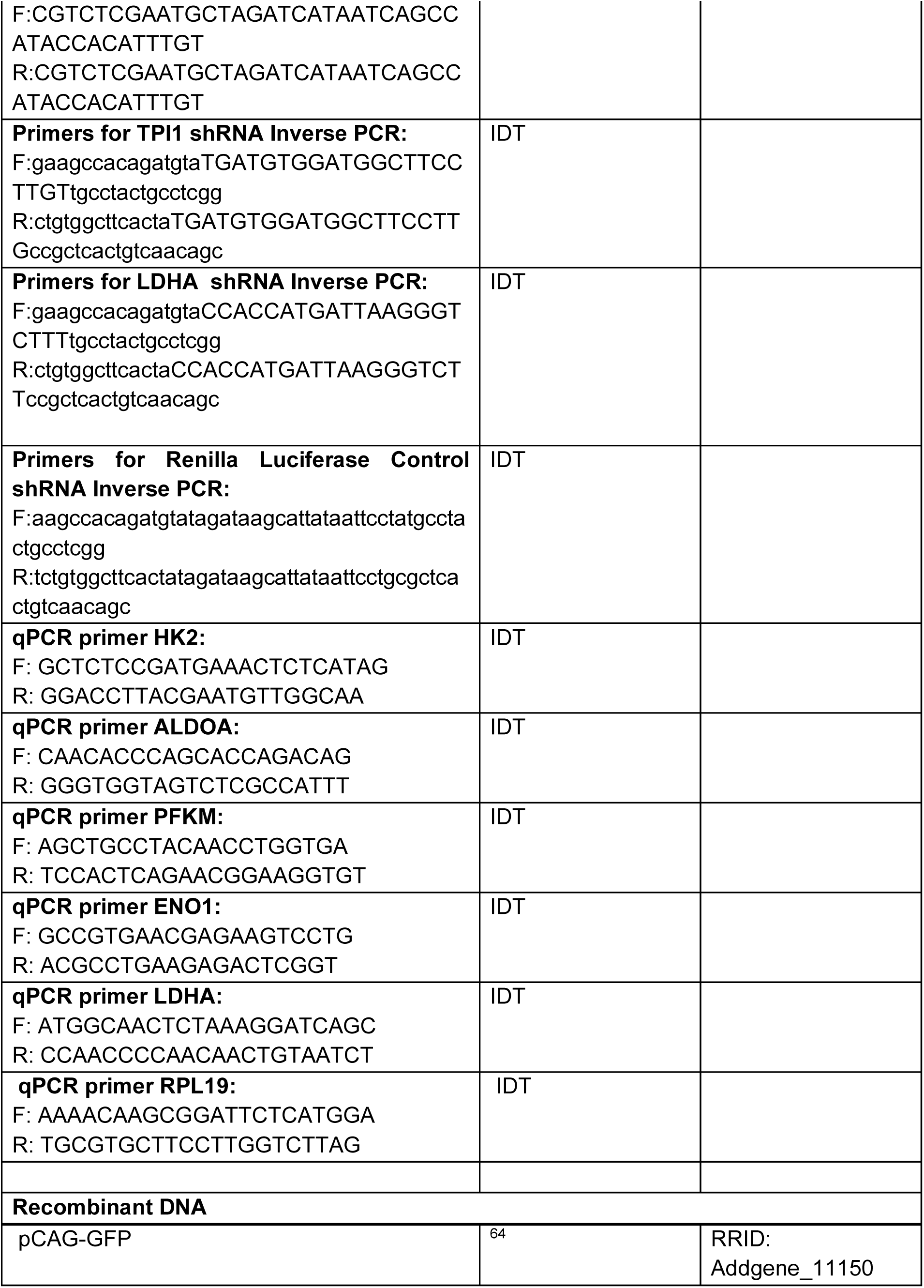
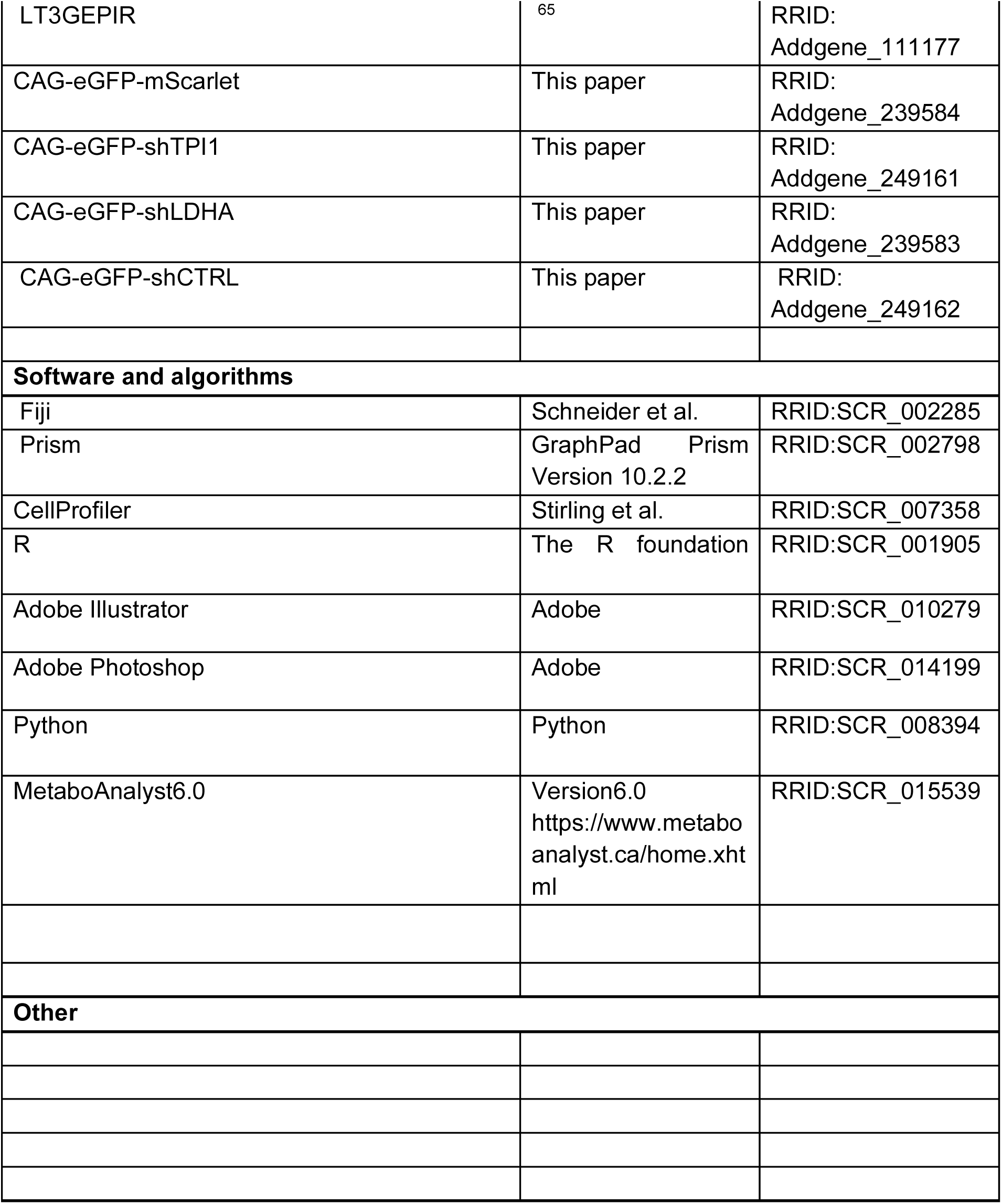

## RESOURCE AVAILABILITY

### LEAD CONTACT

Further information and requests for resources and reagents should be directed to and will be fulfilled by the lead contact, Madeline Andrews.

### MATERIALS AVAILABILITY

All materials generated will be made available by request to the lead contact.

## EXPERIMENTAL MODEL AND STUDY DETAILS

### Pluripotent Stem Cell Lines

Human PSC lines H1/WA01, H21792, 13234, SCTi003-A, and H28126 were used. PSC lines were authenticated prior to differentiation. Every 10 passages, cells are tested for karyotypic abnormalities and validated for pluripotency markers *SOX2*, *NANOG*, and *OCT4*. All cell lines tested negative for mycoplasma.

### Pluripotent stem cell expansion culture

Human induced pluripotent stem cell lines, 13234, H21792, and SCTi003A, and the embryonic stem cell line, WA01 (H1), were expanded on growth factor-reduced Matrigel-coated six well plates. Cells were thawed in mTESR plus (Stem Cell Technologies) containing 10uM of Rock Inhibitor, Y-27632. The media was changed on alternate days. Each line was passaged using ReLeSR (Stem Cell Technologies) and gentle manual lifting when wells reached ∼75% confluency. Cells were split at 1:6 ratio and plated on fresh Matrigel-coated plates or frozen at - 80C in mFresr (Stem Cell Technologies) for 24 hours before being moved to -150C. All lines used in this study were between passage 20-50.

### Regionalized forebrain organoid differentiation protocol

Organoids are differentiated using a regionalized forebrain differentiation protocol ^32,33,66^. Stem cells were dissociated to single cells using Accutase. After dissociation, cells were reconstituted in neural induction media maintaining a density of 10,000 cells per well of 96-well V-bottom ultra-low adhesion plates. Neural induction media is GMEM-based, containing 25mM glucose, and includes 20% Knockout Serum Replacer (KSR), 1X non-essential amino acids, 0.11mg/mL Sodium Pyruvate, 1X Penicillin-Streptomycin, 1X Primocin, 0.1 mM Beta Mercaptoethanol, 5 μM SB431542 and 3 μM IWR1-endo. Cells were treated with 20 μM Rock inhibitor Y-27632 for the first 6 days. After 18 days, organoids were transferred to 6-well low adhesion plates and moved to an orbital shaker rotating at 90 RPM and transitioned into a DMEM/F12-based media containing 17.5mM glucose (rounded in text to 18mM), 1X Glutamax, 1X N2, 1X CD Lipid Concentrate, 1X Penicillin-Streptomycin, and 1X Primocin. After 35 days, organoids were moved into DMEM/F12-based media containing 10% FBS, 5ug/mL Heparin, 1X N2, and 1X Chemically Defined Lipid Concentrate. At 70 days media was additionally supplemented with 1X B27. Organoids were fed every other day throughout culture duration.

### Glucose and oxygen perturbations

For physiological conditions, organoids were aggregated in respective O_2_ conditions (either 20 or 8%) and slowly weaned off high glucose concentrations (25mM GMEM) with ½ concentration (12.5mM) from days 9 - 18, and fully transitioned to 5mM glucose concentrations by day 18 of differentiation. For glucose conditions, starting at Day 9, Standard media #1 was split in a 50:50 ratio with media #1 with basal media containing 5uM glucose. At day 18, during transfer to media #2 and 6 well plates. Media #2 was composed of DMEM/F12, RPMI, or PLASMAX at 5mM glucose. At Day 35, the organoids were transferred to media #3 containing basal media made of DMEM/F12, RPMI, or PLASMAX. Organoids were maintained in Multigas incubators and cultured at either 20% or 8% O_2_ conditions from aggregation day 0.

## METHOD DETAILS

### Single cell capture and RNA sequencing

Organoids were dissociated with a Papain dissociation kit (Worthington) for 60 minutes, spun down at 300g for 5 min, resuspended in PBS + 0.1% BSA, filtered, and counted while verifying single cell dissociation. Single-cell capture was performed using 10X Genomics Chromium Single Cell 3’ v3.1 kit per manufacturer’s instructions. 10,000 cells were targeted for capture and 12 cycles of amplification for each of the cDNA amplification and library amplification were performed. Libraries were sequenced as per manufacturer recommendation on a NovaSeq X Plus flow cell with a minimum read depth of 250 million reads per sample.

### Organoid and tissue fixation and embedding

Organoids were fixed in 4% PFA for 45 mins at 4°C, washed overnight in 1X PBS at 4°C, and then incubated in 30% sucrose in 1X PBS overnight at 4°C. Once samples are saturated they sink in the 30% sucrose solution. Samples were then embedded in cryomolds in a 50:50 solution mix of O.C.T. (Tissue-tek) and 30% sucrose in 1X PBS. Embedded samples were frozen on dry ice and stored at -80°C. Frozen tissue was sectioned into thinly sectioned at 12-16μm using a cryostat (Leica) on glass slides and stored at -80°C.

### Immunohistochemistry

Antigen-retrieval was performed with 1x boiling citric acid-based antigen unmasking solution (Vector Laboratories) for 20 min. Slides were washed 1X with ddH2O and 3X with 1X PBS to remove the citrate solution. Tissue was blocked using a Donkey blocking buffer (DBB) containing 5% donkey serum, 2% gelatin and 0.1% Triton X-100 in 1X PBS (PBS-T) for >30 minutes at RT. Primary antibodies were incubated in DBB overnight at 4°C, washed 3X times with PBS-T, and incubated with AlexaFluor secondary antibodies (ThermoFisher and Jackson Labs) in DBB for 2 hrs at RT. Samples were washed 3X with PBS-T, 1X PBS, mounted, and dried overnight. Parallel sections were used as negative staining controls that were incubated in either primary or secondary, alone. Control slides were processed with antigen retrieval, washed with 1X PBS, and coverslipped like experimental samples. Negative controls were processed to ensure that fluorescence signal on experimental samples is specific. Images were acquired using a Zeiss LSM 710 or Leica SP8 confocal microscope with laser/gain settings consistent across samples/conditions within a stain combination and experiment.

### Spent media collection and analysis

Every two weeks, during routine media changes, 1mL of spent media was isolated from the culture plate. Media was collected from each line of PSCs and derived organoids from weeks 2, 4, 6, 8, 10, and 12 to capture the temporal range of progenitor and neurogenic development. Media from the same condition that was not cultured on organoids, was collected as negative controls. Spent media was spun down to remove cell debris and transferred into low-bind 1.5ml tubes for 80°C storage. After all temporally-collected spent media samples were isolated, samples were thawed on ice, spun down, and each sample loaded in triplicate on a 96 well plate in 100ul volumes. Metabolite composition was analyzed using a YSI 2900 immobilized enzyme calculator to evaluate the concentration of glycolysis-relevant metabolites, glucose and lactate. Values per metabolite were quantified and normalized to the average size of organoid from each cell line per time point.

### Metabolite isolation

Intracellular metabolites were extracted from five organoids per condition. Organoids were washed with 1 mL cold PBS to remove residual media, then quenched with 1 mL 80:20 UHPLC grade methanol:water at -80 °C. Organoids were lysed by sonication at 4°C, then frozen on dry ice for 30 minutes. Samples were vortexed for 30 seconds, then centrifuged at 12,000 rpm for 10 minutes at 4°C. 800 µL of the supernatant was collected and stored on dry ice. 500 uL of 80% methanol was re-added to the cells at -80 °C, followed by repeating the procedures for a second extraction. Cells were re-sonicated and subjected to another freeze-thaw cycle, then centrifuged again with the same settings. 500 uL of the resulting supernatant was added to the initial extract (final volume of 1.3 mL), then the combined extracts were centrifuged again at 12,000 rpm for 10 minutes at 4°C to remove any debris. A final volume of 1.2 mL was collected into 5 mL Eppendorf vials and diluted with UHPLC grade water to achieve a 20% methanol concentration to freeze at -80 °C. Extracts were dried with lyophilization and stored at -80 °C until LC-MS analysis. Following metabolite extraction, the remaining cell pellet was subsequently used for DNA extraction using the DNneasy Blood and Tissue™ kit (Qiagen). Extracted DNA from each sample was measured on a Nanodrop spectrometer and then used to scale metabolite concentrations.

### Untargeted LC-MS Metabolomics

Untargeted metabolomics experiments on organoids was modeled after previous studies^67–73^. LC-MS experiments were performed on a Thermo Vanquish UPLC-Exploris 240 Orbitrap MS instrument. Extracted metabolites from each organoid sample (per line/timepoint/condition) were resuspended in LC-MS grade 40:60 PBS:ACN. Each sample was analyzed twice using an injection volume of 10 µL for negative ionization mode, and 4 µL for positive ionization mode. Chromatographic separation was performed in hydrophilic interaction chromatography (HILIC) mode on a Waters XBridge BEH Amide column (150 x 2.1 mm, 2.5 µm particle size, Waters Corporation). The flow rate was 0.3 mL/min with an auto-sampler temperature of 4 °C and the column compartment at a temperature of 40 °C. The mobile phase was composed of Solvents A (10 mM ammonium acetate, 10 mM ammonium hydroxide in 95% H_2_O/5% ACN) and B (10 mM ammonium acetate, 10 mM ammonium hydroxide in 95% ACN/5% H_2_O). After the initial 1 min isocratic elution of 90% B, the percentage of Solvent B decreased to 40% at t=11 min. The composition of Solvent B maintained at 40% for 4 min (t=15 min), and then the percentage of B gradually went back to 90%, to prepare for the next injection. Untargeted mass spectra were collected from 70 to 1050 m/z using an electrospray ionization source. A Quality control (QC) sample was also prepared as the mixture of each organoid sample, and it was measured once per 10 study samples to ensure measurement reproducibility.

### shRNA cloning

eGFP, mScarlet, and a backbone with the CAG promoter were amplified with primers containing BsmbI golden gate assembly overhangs using Q5® High-Fidelity DNA Polymerase (New England Biolabs) and then gel extracted. The CAG backbone inverse PCR reaction was digested with DpnI, digesting the original template. Golden Gate Assembly was performed overnight using BsmbI and T4 DNA ligase (New England Biolabs) with 50x cycles of 42°C for 2 minutes and 16°C for 5 minutes followed by a single 60°C 5-minute hold. The golden gate reaction was transformed into One Shot™ MAX Efficiency™ DH5α-T1R Competent Cells (Invitrogen) and outgrown at 30^C^ for 16 hours. Liquid cultures were made using single colonies in LB broth plus 50 mg/mL kanamycin, then miniprepped according to the Monarch® Plasmid Miniprep Kit (New England Biolabs). A differential digest was performed using restriction enzymes BamHI and EcoRV and run on a gel for 40 minutes at 100V. Minipreps with expected results were sequenced via Plasmidsaurus. CAG-eGFP-mScarlet was linearized with XhoI and MfeI, removing the mScarlet fragment. The linearized backbone and insert were recircularized using NEBuilder® HiFi DNA Assembly and transformed, miniprepped, digested with EcoRI/BamHI, and sent for sequencing as described above. shRNAs for TPI1, LDHA, and a control targeting Renilla Luciferase were constructed through inverse PCR using Q5® High-Fidelity DNA Polymerase and processed similarly to confirm the expected sequence. shRNA plasmids were maxiprepped according to the EndoFree® Plasmid Maxi Kit (QIAGEN) prior to electroporation. All primers used were ordered from Integrated DNA Technologies and all *in silico* design was performed using Benchling. All restriction enzymes were from New England Biolabs. mRNA target sites for shRNAs were derived from splashRNA and Millipore Sigma MISSION® RNAi ^74^.

### Organoid electroporation

Control (Renilla Luciferase), TPI1, and LDHA shRNAs were electroporated in week 6 forebrain organoids. Six organoids per condition were placed in a 400mm cuvette and bathed in 100 µl media containing 1ug/ul plasmid for 10mins. After incubation, using a ECM 830 BTX electroporator with an attached safety dome, organoids were electroporated at a voltage of 200V for a duration of five milliseconds and frequency of three pulses. Organoids were floated up from the cuvette by increasing the media volume to 200ul. The media and organoids were returned to six well plates and cultured under standard conditions for one week before processing.

### Organoid drug experiments

Organoids were differentiated for 6 weeks, as before. For PFK and LDHA perturbations, stock solutions of each inhibitor were prepped in DMSO at 10mM and 20µM, respectively. Inhibitors or DMSO were diluted 1:1000 in 3mL media to achieve final concentrations of 10µM and 20nM for PFK and LDHA, respectively. Organoids were treated for 48 hours, then collected for IHC and q-RT-PCR. For lactate experiments, 3mL of standard (DMEM/F2 with 18mM glucose) media with or without 10mM sodium-L-lactate (Sigma). After 48 hours, organoids were collected and processed for IHC.

### Western Blot

Organoids were lysed in RIPA buffer supplemented with Halt protease inhibitor (100X) and EDTA (0.5M). After a 30-minute incubation on ice, lysates were centrifuged at 20,000xg for 30 minutes at 4°C. Protein samples were prepared in Bolt LDS sample buffer with Bolt sample reducing agent and nuclease free water, heated at 70°C for 10 minutes, and separated by electrophoresis on 4-12% Bis-Tris gels at 200V for 20 minutes. Proteins were transferred to nitrocellulose membranes using an iBlot3 system, blocked in 5% non-fat milk for 60 minutes, and incubated with primary antibodies in 5% BSA overnight at 4°C. Membranes were washed three times for 10 minutes each in 1X TBST, followed by secondary antibody incubation in 1% BSA for 1 hour at room temperature. Membranes were then washed 6X for 5 minutes each in 1X TBST, developed using SuperSignal West Pico or SuperSignal West Atto substrate, and imaged on an Invitrogen iBright Imaging System for quantitative analysis via the iBright™ Analysis Software.

### q-RT-PCR

3 organoids were collected at week 4, 6, 8, 12, 15/16. The organoids were lysed by mechanical shearing in RLT buffer and then stored at -80C. RNA was extracted using Qiagen RNeasy Plus Micro Kit (Cat# 74134), RNA concentration and purity was determined using a NanoDrop spectrophotometer, and then stored at -80C. cDNA was prepared using SuperScript VILO cDNA Synthesis Kit (Cat# 11754050). qPCR reactions was performed using iTaq Universal SYBR Green Supermix (Cat# 1725121) and primers listed below. Each experiment was assessed in technical triplicate, with replication across multiple cell lines. Gene expression fold changes were determined using the 2-method, with RPL19 serving as the normalization control.

## QUANTIFICATION AND STATISTICAL ANALYSIS

### Quantification and statistical analysis of immunohistochemistry

A CellProfiler-based (version 4.2.8)^75^ pipeline was developed to analyze the expression of markers of interest in organoid sections. Multi-channel LSM files were first processed in FIJI to generate maximum intensity z-projections. Individual channels were separated and exported as single channel images for subsequent analysis. Cell identification and marker quantification were performed using CellProfiler. Cell markers were identified using a two-class Otsu thresholding method or minimum cross-entropy thresholding. The pipeline was optimized to identify positive cells based on fluorescence intensity thresholds and cellular morphology. For all markers, positive cells were filtered by diameter and brightness and normalized to DAPI counts. Cell counting was automated using the pipeline parameters calibrated to match manual counts. Six images per condition were analyzed. For statistical analysis, data normality was assessed using the Shapiro-Wilk test in GraphPad Prism. For comparisons between two different conditions, an unpaired t-test was applied when both groups passed the normality test (p>0.05). If either group failed to meet normality assumptions, non-parametric Mann-Whitney U test was used. For experiments with more than two conditions being compared, if normally distributed, an ordinary one-way ANOVA was used and if not normally distributed, the non-parametric Kruskal-Wallis test was used.

### Metabolite Data Identification

Identification of MS spectra peaks was performed using in-house chemical standards (∼600 aqueous metabolites). The resulting MS spectra were compared against publicly available libraries: HMDB library, Lipidmap database, METLIN database; and commercial databases: mzCloud, Metabolika, and ChemSpider. The absolute intensity threshold for the MS data extraction was 1,000, and the mass accuracy limit was set to 5 ppm. Peaks were identified and labeled using available data for retention time (RT), exact mass (MS), MS/MS fragmentation pattern, and isotopic pattern. Thermo Compound Discoverer 3.3 software was used for aqueous metabolomics data processing. Untargeted data was processed for peak picking, alignment, and normalization. Inclusion criteria for analysis included signals/peaks with coefficient of variation (CV) < 20% across QC pools and the signals showing up in >80% of all the samples.

### Metabolite Data Analysis

The resulting data from each sample was normalized to its corresponding DNA concentration. PQN normalization was performed using median scaling, the total signal from each sample (i.e. sum of all metabolites captured), to estimate a normalization coefficient that reduces technical variability between each run using an established method ^76^. The data was then filtered to remove exogenous and non-organic compounds then Log10 transformed to directly compare metabolite abundances.

### Z-score & Heatmap visualization

An average z score was computed by calculating the average metabolite abundance for each metabolite across replicates per condition. A z-score function was performed on the averages to compare increases/decreases between conditions. The changes in Z-score per metabolite were visualized for heatmap comparison. Unpaired t tests with individual variances were used to identify significantly different metabolites between each group. We then computed FDR adjusted p-values to correct for multiple comparisons.

### Metabolite Enrichment Pathway analysis

Metabolite pathway analysis was performed using **MetaboAnalyst6.0** ^77^. Log fold change analysis was used to generate lists of metabolites with a log2FC ≥ 0.25 for each condition. Metabolite enrichment analysis with KEGG library was performed for each list of metabolites to identify significantly enriched pathways in each condition using an FDR <= 0.1.

### scRNAseq pre-processing and QC

Cellranger (v8.0.1, 10x Genomics) was used for data pre-processing. Barcode correction and reference genome alignment (GRCh38) were performed using cellranger count with default parameters on fastq files for each organoid sample. Python (v3.12.7) was used to subset the original primary dataset before being piped into Seurat v5 for analysis. For each organoid sample, count matrices were piped into Seurat v5 ^78^, in which cells with fewer than 200 genes and genes present within less than 3 cells were removed from each of the 36 organoid samples. Ensembl IDs not converted to gene symbols during cellranger count alignment were removed. Each individual sample was further filtered using cutoff values (S Table 3) that were visually determined using scatter plots of mitochondrial counts vs total counts. Mitochondrial counts were capped at 10% for each sample.

### sc-RNA-seq processing and clustering

After each sample was processed, all 36 samples were combined using the merge function in Seurat and labeled with the appropriate cell ID. The merged object (248,661 cells) was used for all downstream analysis. Raw counts from individual samples were normalized by SCTransform^79^. Subsequently, all the organoid data were integrated using the RPCA methods in the R package Seurat v5 ^80^. Principal component analysis was performed using RunPCA, and significant principal components were identified. In the space of these significant principal components, the k = 10 nearest neighbors were determined and identified using the FindNeighbors function. If any clusters contained only one cell, the process was repeated with k = 11 and upwards until no clusters contained only one cell.

### Annotations

Cell type annotations of clusters were performed by comparison to previously annotated cell types ^29^, and when a repository of substantial matching was not available, a combination of literature-based annotation of layer or maturation stage identity and output from the Seurat function FindAllMarkers was used. UMAP embedding was computed for visualization. Feature plots showing canonical cell type markers were also used to aid in annotation.

### Differential Gene Expression & Functional Enrichment

Differential gene expression was performed using DESeq2 v1.46.0 ^81^ on pseudobulk data relating to each overarching cell type across experimental conditions. Each cluster of cells from the original clustering and annotation was identified by the overarching cell type (i.e. ‘oRG’) and merged into a single cluster consisting of that cell type until all original clusters were represented by 11 distinct cell types. The raw RNA counts from each newly combined category were then shuttled into the DESeq2 pipeline to assess differential expression of genes across experimental conditions using default parameters. After the lists of DEGs were created, the genes passing a threshold of adjusted p value < 0.05 (Benjamini-Hochberg (BH) method) and log2 fold change <0.2 were visualized using EnhancedVolcano v1.24.0 ^82^. Functional enrichment was assessed using clusterProfiler v4.16.0 ^83^ to convert the DEG symbols to Entrez IDs with subsequent biological process gene ontology (GO) term enrichment using the enrichGO function with default parameters. This same pipeline was applied to all cells together across experimental conditions as well (i.e. the pseudobulk of all cells at week 6 tested against all cells at week 10 with respect to the media and oxygen conditions).

### Hypergeometric Test

Hypergeometric distribution analysis was conducted to find the number and identity of statistically significant (adjusted p value < 0.05 using BH) shared genes between sets of DEGs found using DESeq2. The analysis was conducted using the Hypergeometric function phyper within the r package stats v4.5.0. The parameters of this function were defined and arranged to calculate the probability of observing the number of shared genes where 1 is subtracted from the number of shared genes and the lower.tail parameter is set to false. The names of the genes found in each intersection that were also statistically significant (adjusted p value < 0.05 using BH) were assessed for biological process GO term enrichment in the same way as described above. This same workflow was also applied to the genes identified as unique in each comparison.

### Determining the flux through metabolic pathways using METAFlux

Flux balance analysis was computed from three biological replicate (iPSC line) pseudobulk samples from the scRNA-seq data using METAFlux^43^. The combined annotated cell types from the single cell analysis pipeline were split into three equally sized and aggregated into unique pseudobulk RNA-seq replicate populations using the “createDataPartition” function from the caret R package (v7.0-1). Summed pseudobulk counts files were generated using the AggregateExpression function from the seurat pipeline and normalized using the vst function in DESeq2 (v1.30.1) for each unique cell type within each timepoint. Flux analysis was then performed using the METAFlux bulk RNA-seq pipeline using the default “cell_medium” nutrient microenvironment and human-GEM files. Flux scores were transformed using cubic root normalization prior to pathway level activity quantification. Cell line, timepoint, oxygen, and media information were recorded for each sample and used in downstream pathway comparisons. Significant differences in flux between control and test samples of each condition were determined using two-tailed Student’s t-tests and the magnitude of the effect represented using log_2_(FC) of the median flux between triplicates. Metabolic pathways were considered significant for a condition if the |log_2_(FC)| ≥ 1.0 with a p-value ≤ 0.05. Heatmaps used to represent the data and were generated using the seaborn Python data visualization package (v0.13).

## ACKNOWLEDGMENTS

The authors thank Drs. Aparna Bhaduri, Caroline Pearson, Miyeko Mana, and Connor Dolan, and members of the M.G.A. laboratory for useful discussion. We thank Dr. Joy Blain and Anna Engelbrektson from the ASU Genomics Core for their support with library preparation. This study was supported by NIH award R00MH125329, Arizona Biomedical Research Centre grant #RFGA2022-010-16 to M.G.A, and ASU-Mayo Seed Grant to C.F. & M.G.A.

## AUTHOR CONTRIBUTIONS

Conceptualization: T.P., A.S.M.M, C.F., M.G.A.; Investigation: T.P., A.M., S.C., A.S.M.M., G.B., J.C., F.F., T.N., A.M.S., L.L., M.G.A; Data Curation: A.S.M.M, T.P., C.P.; Methodology: A.M., S.C., T.P., A.S.M.M. B.B., F.F., M.G.A.; Formal analysis: A.S.M.M, T.P., A.M., S.C., S.W., C.P.; Writing– original draft: T.P., A.S.M.M, M.G.A.; Writing – review & editing: T.P, A.S.M.M, S.C., M.G.A.; Funding acquisition: M.G.A., C.F.; Resources: M.G.A., B.B., H.G., C.P., C.F.; Supervision: M.G.A., B.B., C.P., H.G., C.F.

---

**S Figure 1:**
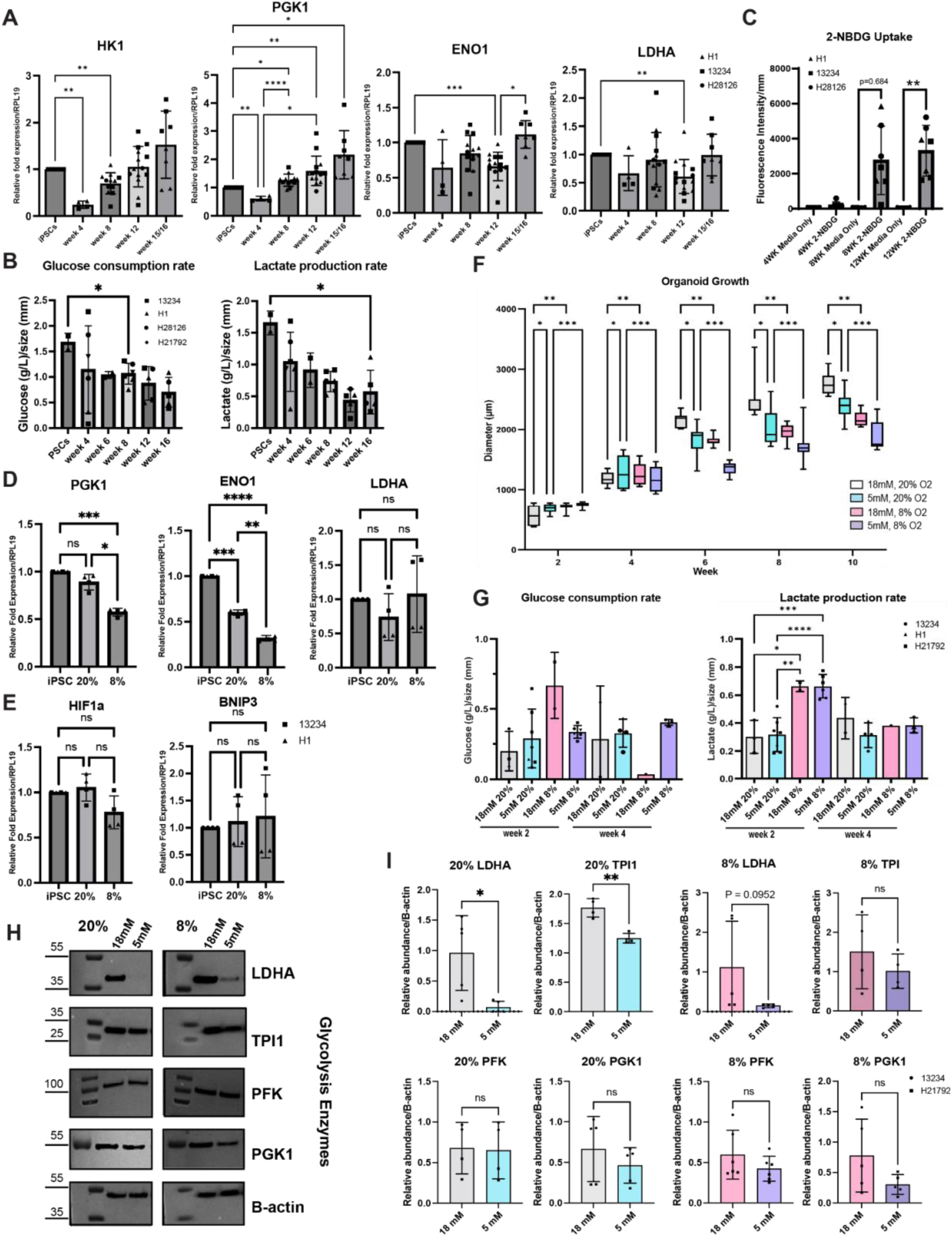
Organoids cultured under standard conditions have unclear metabolic transitions. **A)** q-RT-PCR on glycolysis genes was performed on iPSCs and organoids from the same lines until 16 weeks of differentiation (HK1: One-way ANOVA, iPSC vs wk4: **p<0.0013, iPSC vs wk8: **p<0.0058; ENO1: One-way ANOVA, iPSC vs wk12: ***p<0.0002, wk12 vs wk15/16: *p<0.0154; LDHA: iPSCs vs wk12: **p<0.0025; PGK1: iPSC vs wk4: **p<0.0097, iPSC vs wk8: *p<0.0112, iPSC vs wk12: **p<0.0072, iPSC vs wk15/16: *p<0.0366; wk4 vs wk8: ****p<0.0001; wk4 vs wk12: *p<0.0279). n=4 organoids pooled per differentiation batch from n=3 iPSC lines are collected for q-RT-PCR repeated n=2 times each per line. Data are mean ± SD. **B)** Spent media analysis. Glucose consumption minimally changed over time under baseline culture conditions (Ordinary one-way ANOVA with Tukey’s multiple comparisons, PSC vs wk8: *p<0.0211). Lactate production gradually decreased without significant change until gliogenic periods (Ordinary one-way ANOVA with Tukey’s multiple comparisons, PSC vs wk16: *0.0211). Spent media collected from organoids from n=4 lines across time points. Data are represented as mean ± SD. **C)** Uptake of 2-NBDG in organoids increases during neurogenic and gliogenic stages (Ordinary one-way ANOVA, Wk8 media vs Wk8 2-NBDG: p=0.0684; Wk12 media vs Wk12: **p<0.0069). n=3 organoids per differentiation batch each from n=3 iPSC lines. Data are represented as mean ± SD. **D)** q-RT-PCR was performed in organoids cultured under 20 vs 8% O_2_ conditions. Under 8% conditions, expression of glycolysis genes *PGK1* and *ENO1* decreased (Ordinary one-way ANOVA with Tukey’s multiple comparisons, PGK1: iPSCs vs 8% O_2_ : ***p<0.0005; 20% vs 8% O_2_: *p<0.0101; ENO1: iPSCs vs 20% O_2_ : ***p<0.0002; iPSCs vs 8% O_2_ : ****p<0.0001; 20% vs 8% O_2_: **p<0.0032). n=4 organoids pooled per differentiation batch from n=3 iPSC lines are collected for q-RT-PCR repeated n=2 times per line. Data are represented as mean ± SD. **E)** q-RT-PCR was performed on organoids cultured under 20 vs 8% O_2_ conditions. Under 8% conditions, expression of hypoxia genes *HIF1a* and *BNIP3* did not change. n=4 organoids pooled per differentiation batch from n=3 iPSC lines are collected for q-RT-PCR and repeated n=2 times each per line. Data are represented as mean ± SD. **F)** Organoid size changes over time and under different conditions (Two-way ANOVA with multiple comparisons: 18mM 20% vs 5mM 20%: *p=0.0229, 18mM 20% vs 18mM 8%:**p=0.0012, 5mM 20% vs 5mM 8%: ***p=0.0001, n>7 organoids/timepoint from n=3 PSC lines). Data are represented as mean ± SD. **G)** Glucose consumption per conditions for weeks 2-4 shows most significant increases in 8% O_2_ conditions (Ordinary one-way ANOVA with Tukey’s multiple comparisons, n>3 spent media collections per timepoint from n=3 lines). Lactate production for week 2-4 with most significant increase in 8% O_2_ conditions at week 2 (Ordinary one-way ANOVA with Tukey’s multiple comparisons: Week 2: 18mM 20% vs 18mM 8%: *p<0.0267; 18mM 20% vs 5mM 8%: ***p<0.0006; 5mM 20% vs 18mM 8%: **p<0.0093; 5mM 20% vs 5mM 8%: ****p<0.0001; n>3 spent media collections per timepoint from n=3 lines). Datapoints are metabolite concentration values/line ± SD. **H-I)** Western blot analysis of glucose metabolism enzymes. Under 20% O_2_ conditions, LDHA and TPI1 decreased after treatment with 5mM glucose, but there were no significant changes in other glucose metabolism proteins. (20% LDHA: Shapiro-Wilk normality test: *p<0.027, Mann-Whitney test: *p<0.0159; 20% TPI1: Shapiro-Wilk normality test: p=0.143, unpaired t test: **p<0.0011). n=4 organoids pooled per differentiation batch from n=2 iPSC lines are collected for western blot and repeated n=2 times each per line. Data are represented as mean ± SD.

**S Figure 2.**
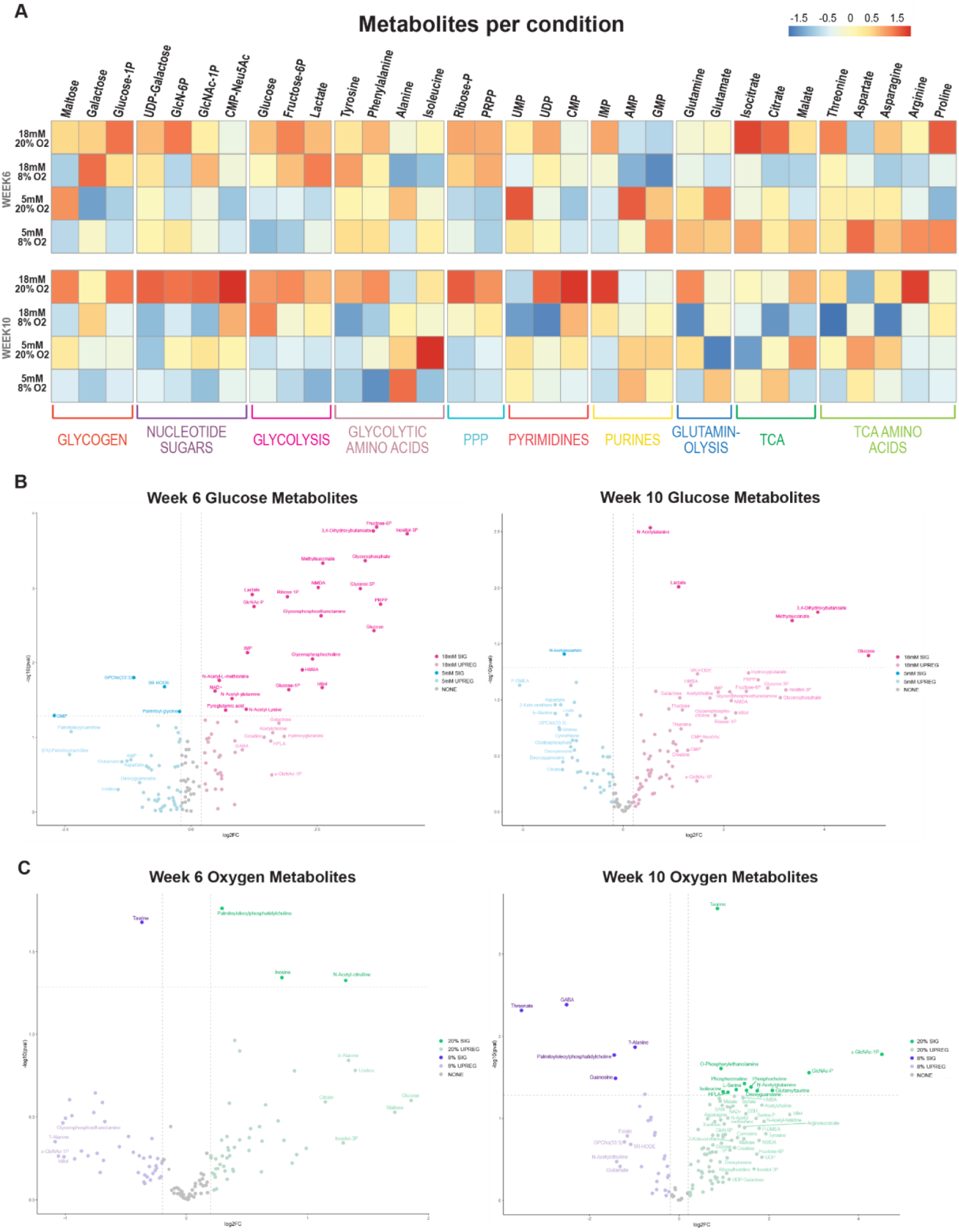
Glucose and oxygen conditions impact metabolite abundance. **A)** Heatmap summarizes changes in metabolite composition per condition across time. Metabolites were extracted from n=5 organoids pooled per differentiation batch from n=3 iPSC lines/condition. **B)** 18mM glucose conditions had more metabolites significantly upregulated compared to 5mM glucose, particularly glycolysis regulators, such as glucose, lactate, and Fructose-6P, at both 6 and 10 weeks of differentiation. **C)** Few metabolites differed between O_2_ conditions at week 6, but at week 10, 8% O_2_ had increase in GABA, Threonate, L-Alanine and 20% O_2_ had increased Taurin, GlcNAc-P, O-Phosphorylethanolamine. n=5 organoids pooled per differentiation batch from n=3 iPSC lines/condition were collected for metabolite extraction.

**S Figure 3.**
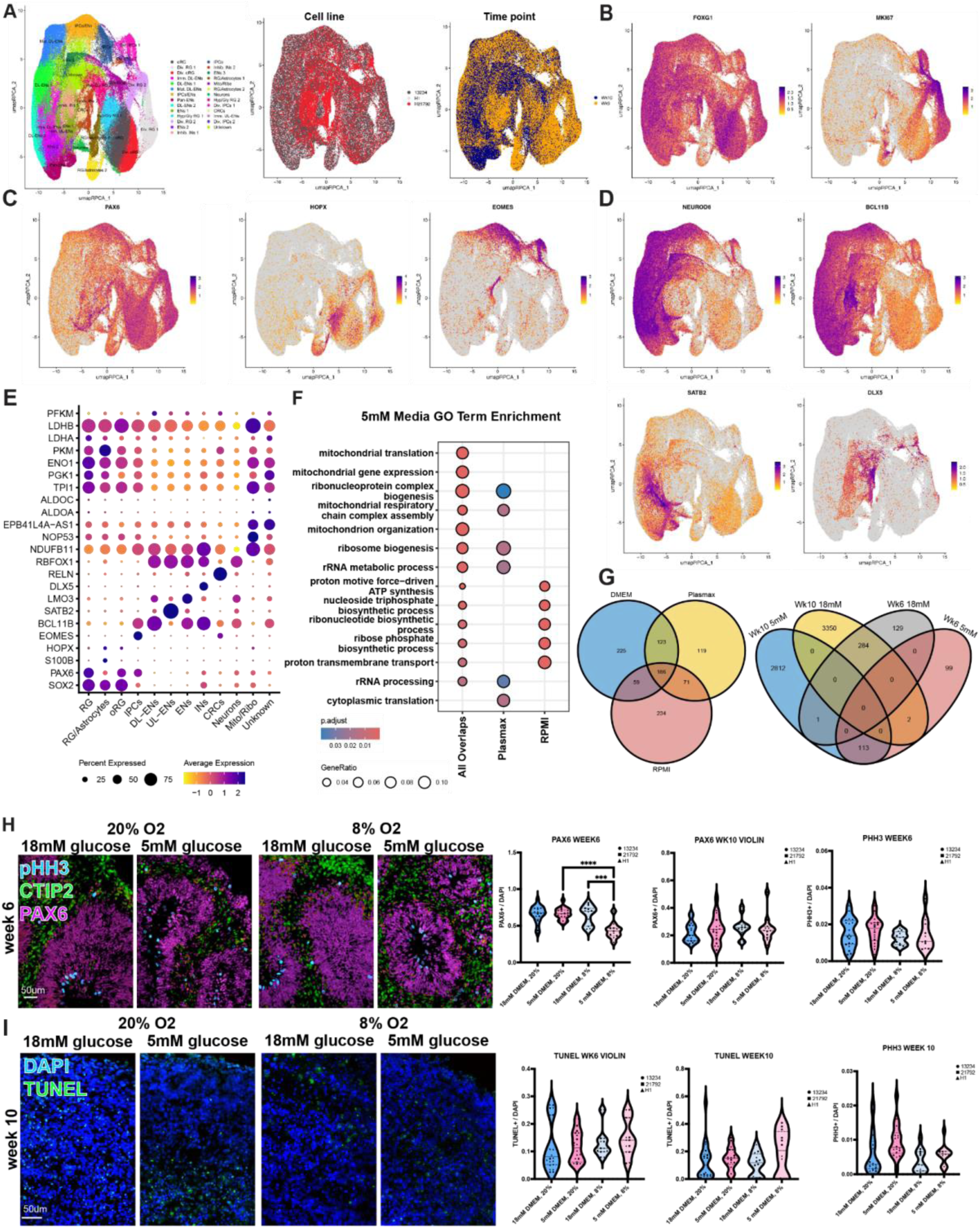
Physiologic glucose and oxygen conditions have minimal impact on proliferation, lineage shifts, & cell death in organoids. **A)** UMAPs of collected organoid data by more granular cluster annotation, iPSC line, and timepoint of collection. **B-D)** Feature plots of cortical (*FOXG1*), progenitor (*KI67, PAX6, HOPX, EOMES*), and neuronal marker (*NEUROD6, BCL11B, SATB2, DLX5*) genes. **E)** Dotplot indicates marker expression and percent of cells each gene is expressed across clusters. Canonical cell type marker genes, top DEGs for each cell type, and glycolysis genes are shown. **F)** GO term enrichment dotplot of significantly enriched terms found in all overlapped 5mM media DEGs from all cells across timepoints (DMEM/F12, Plasmax, RPMI) compared against the unique DEGs to each media (Plasmax, RPMI). DMEM/F12 had no significantly enriched terms. **G)** Venn diagrams showing the overlapping DEGs from all cells regardless of timepoint between medias (DMEM/F12, Plasmax, RPMI; left diagram) and the DEGs from all cells across glucose concentration (18mM , 5mM) and timepoint (wk6, wk10) (right diagram). **H)** Organoids cultured under different glucose and oxygen conditions had minimal impact on PAX6+ RG without impacting pHH3+ proliferation. At week 6 there was a small decrease at week 6 under 8% O_2_ conditions (One way ANOVA, PAX6 week6: 5mM 20% vs 5mM 8%: ****p<0.0001, 18mM 8% vs 5mM 8%: ***p<0.0001). **I)** Organoids cultured under different glucose and oxygen conditions did not increase the number TUNEL+ dying cells. n=4 organoids per differentiation batch each from n=3 iPSC lines/condition/timepoint. Data are represented as mean ± SD.

**S Figure 4:**
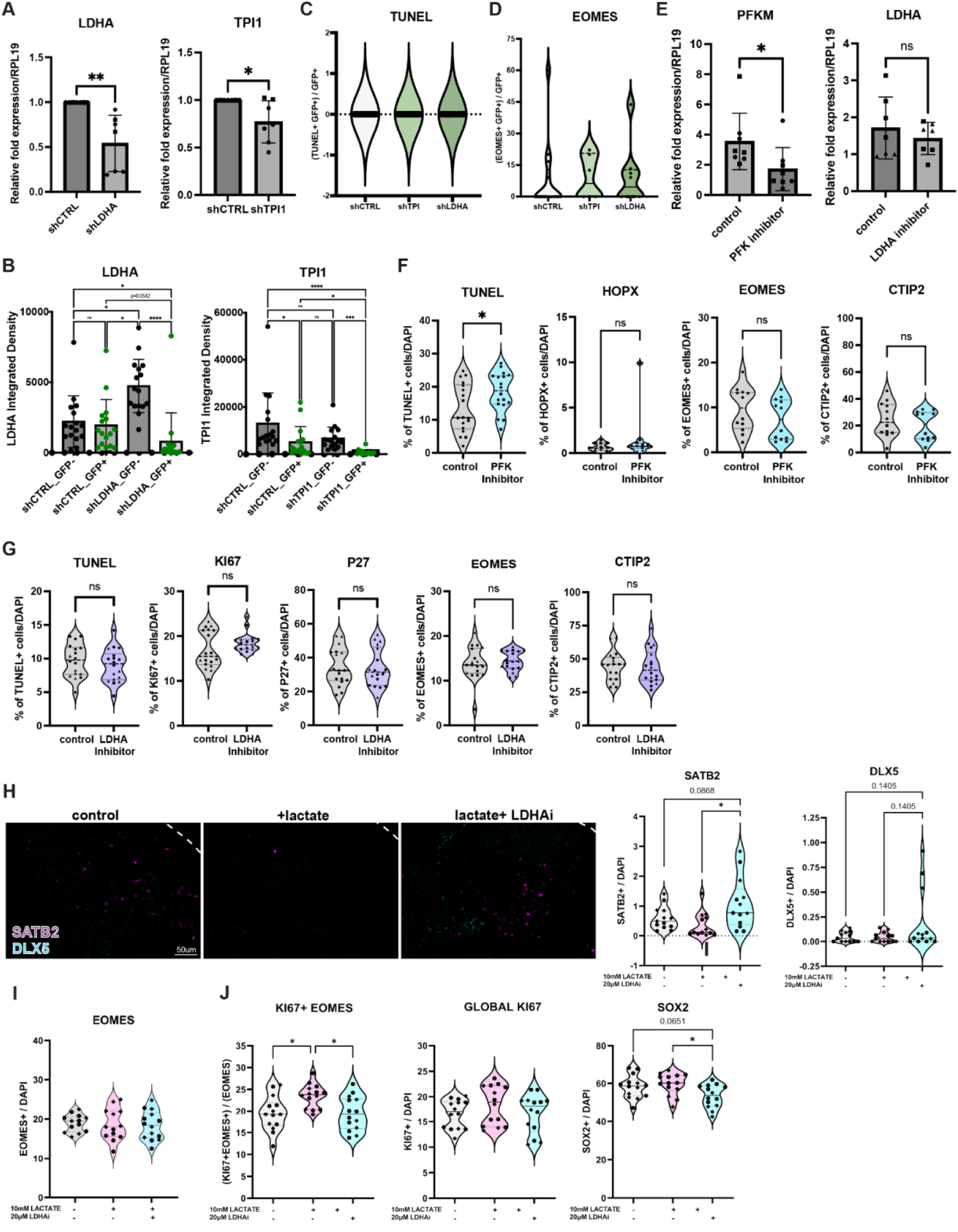
Changes in glycolysis have cell-specific impact during neurodevelopment. **A)** Electroporation of LDHA and TPI1 shRNAs decreased levels of respective RNA expression by q-RT-PCR (LDHA: Welch’s t-test: **p<0.0083; TPI1: Welch’s t-test: *p<0.0337). **B)** Electroporation of LDHA and TPI1 shRNAs result in decreased abundance of targeted protein in electroporated GFP+ cells with knockdown (One way ANOVA with multiple comparisons, shcontrol GFP+ vs shLDHA GFP+: p=0.0542; shLDHA GFP- vs shLDHA GFP+: p<0.0001; shcontrol GFP- vs shLDHA GFP+: *p<0.0157; shcontrol GFP+ vs shTPI1 GFP+: *p<0.0397; shTPI1 GFP- vs shTPI1 GFP+: ***p<0.0001). **C-D)** There are no significant changes in TUNEL+ or EOMES+ cells after electroporation of shTP1 or LDHA (One way ANOVA, EOMES: shTPI1: p=0.9885; shLDHA: 0.9672). **F)** Week 6 organoids treated with PFK inhibitor, PFK15, had decreased relative PFK expression via q-RT-PCR compared to controls (Unpaired student’s t-test: *p<0.0464). Treatment with LDHA inhibitor, GSK2837808A, did not impact LDHA relative expression (Unpaired student’s t-test: p=0.676) **G)** PFK inhibition resulted in a small increase in TUNEL+ dying (Unpaired student’s t-test: *p<0.0282), without change to HOPX+ oRG (Shapiro-Wilk normality test: p<0.0001; Mann-Whitney test: p=0.3186), EOMES+ IPC (unpaired t test: p=0.2621) or CTIP2+ DL-EN (unpaired t test: p=0.3612). **H)** LDHA inhibition did not impact number TUNEL+ dying (unpaired t test: p=0.3752), KI67+ proliferating cells (Shapiro-Wilk normality test: p=0.0134; Mann-Whitney test: p=0.161), p27+ differentiating cells (unpaired t test: p=0.8708), EOMES+ IPCs (unpaired t test, p=0.5612), or CTIP2+ DL-EN (unpaired t test, p=0.8819). n=4 organoids per differentiation batch each from n=3 iPSC lines/condition. **I)** Lactate alone did not significantly affect the proportion of SATB2+ UL-ENs (control vs lactate: p= 0.3220) or DLX5+ IN (control vs lactate: p= 0.1502) at week 6, while the presence of LDHAi led to a significant increase in SATB2+ UL-ENs (lactate vs LDHA inhibitor: *p= 0.0116, control vs lactate+LDHAi: p= 0.0868) and trending increase in DLX5+ interneurons (lactate vs LDHA inhibitor: p = 0.1405, control vs lactate+LDHAi: p= 0.1405). **J)** Lactate with and without LDHAi had no impact on EOMES+ cells (One way ANOVA: lactate: p=0.9542, lactate+LDHAi: p=0.9590). Lactate significantly increased KI67+EOMES+ IPCs, which normalized in the presence of LDHAi (control vs lactate: *p=0.0145; lactate vs LDHA inhibitor: *p=0.0145), but did not impact global KI67+ cells (control vs lactate: p=0.4998 ; lactate vs LDHA inhibitor: p=0.4998, control vs lactate+LDHAi: p=0.9610). The proportion of SOX2+ cells was unaffected by lactate but significantly decreased with addition of LDHAi (control vs lactate: p=0.5497; lactate vs LDHA inhibitor: *p=0.0228, control vs lactate+LDHAi: p=0.0651).

## References

1. Molnár, Z., Clowry, G.J., Šestan, N., Alzu’bi, A., Bakken, T., Hevner, R.F., Hüppi, P.S., Kostović, I., Rakic, P., Anton, E.S., et al. (2019). New insights into the development of the human cerebral cortex. J. Anat. 235, 432–451.

2. Geschwind, D.H., and Rakic, P. (2013). Cortical evolution: judge the brain by its cover. Neuron 80, 633–647.

3. Eze, U.C., Bhaduri, A., Haeussler, M., Nowakowski, T.J., and Kriegstein, A.R. (2021). Single-cell atlas of early human brain development highlights heterogeneity of human neuroepithelial cells and early radial glia. Nat. Neurosci. 24, 584–594.

4. Braun, E., Danan-Gotthold, M., Borm, L.E., Lee, K.W., Vinsland, E., Lönnerberg, P., Hu, L., Li, X., He, X., Andrusivová, Ž., et al. (2023). Comprehensive cell atlas of the first-trimester developing human brain. Science 382, eadf1226.

5. Ziffra, R.S., Kim, C.N., Ross, J.M., Wilfert, A., Turner, T.N., Haeussler, M., Casella, A.M., Przytycki, P.F., Keough, K.C., Shin, D., et al. (2021). Single-cell epigenomics reveals mechanisms of human cortical development. Nature 598, 205–213.

6. Zhu, K., Bendl, J., Rahman, S., Vicari, J.M., Coleman, C., Clarence, T., Latouche, O., Tsankova, N.M., Li, A., Brennand, K.J., et al. (2023). Multi-omic profiling of the developing human cerebral cortex at the single-cell level. Sci. Adv. 9, eadg3754.

7. Nano, P.R., Fazzari, E., Azizad, D., Martija, A., Nguyen, C.V., Wang, S., Giang, V., Kan, R.L., Yoo, J., Wick, B., et al. (2025). Integrated analysis of molecular atlases unveils modules driving developmental cell subtype specification in the human cortex. Nat. Neurosci. 28, 949–963.

8. Dong, X., Zhang, Q., Yu, X., Wang, D., Ma, J., Ma, J., and Shi, S.-H. (2022). Metabolic lactate production coordinates vasculature development and progenitor behavior in the developing mouse neocortex. Nat. Neurosci. 25, 865–875.

9. Bakare, A.B., Lesnefsky, E.J., and Iyer, S. (2021). Leigh syndrome: A tale of two genomes. Front. Physiol. 12, 693734.

10. Klepper, J., Akman, C., Armeno, M., Auvin, S., Cervenka, M., Cross, H.J., De Giorgis, V., Della Marina, A., Engelstad, K., Heussinger, N., et al. (2020). Glut1 Deficiency Syndrome (Glut1DS): State of the art in 2020 and recommendations of the international Glut1DS study group. Epilepsia Open 5, 354–365.

11. Zhao, Y., Fung, C., Shin, D., Shin, B.-C., Thamotharan, S., Sankar, R., Ehninger, D., Silva, A., and Devaskar, S.U. (2010). Neuronal glucose transporter isoform 3 deficient mice demonstrate features of autism spectrum disorders. Mol. Psychiatry 15, 286–299.

12. Merker, S., Reif, A., Ziegler, G.C., Weber, H., Mayer, U., Ehlis, A.-C., Conzelmann, A., Johansson, S., Müller-Reible, C., Nanda, I., et al. (2017). SLC2A3 single-nucleotide polymorphism and duplication influence cognitive processing and population-specific risk for attention-deficit/hyperactivity disorder. J. Child Psychol. Psychiatry 58, 798–809.

13. Modabbernia, A., Mollon, J., Boffetta, P., and Reichenberg, A. (2016). Impaired gas exchange at birth and risk of intellectual disability and autism: A meta-analysis. J. Autism Dev. Disord. 46, 1847–1859.

14. Scafidi, J., and Gallo, V. (2008). New concepts in perinatal hypoxia ischemia encephalopathy. Curr. Neurol. Neurosci. Rep. 8, 130–138.

15. Prigione, A., Fauler, B., Lurz, R., Lehrach, H., and Adjaye, J. (2010). The senescence-related mitochondrial/oxidative stress pathway is repressed in human induced pluripotent stem cells. Stem Cells 28, 721–733.

16. Zhang, J., Khvorostov, I., Hong, J.S., Oktay, Y., Vergnes, L., Nuebel, E., Wahjudi, P.N., Setoguchi, K., Wang, G., Do, A., et al. (2011). UCP2 regulates energy metabolism and differentiation potential of human pluripotent stem cells: UCP2 regulates hPSC metabolism and differentiation. EMBO J. 30, 4860–4873.

17. Bélanger, M., Allaman, I., and Magistretti, P.J. (2011). Brain energy metabolism: Focus on astrocyte-neuron metabolic cooperation. Cell Metab. 14, 724–738.

18. Hall, C.N., Klein-Flügge, M.C., Howarth, C., and Attwell, D. (2012). Oxidative phosphorylation, not glycolysis, powers presynaptic and postsynaptic mechanisms underlying brain information processing. J. Neurosci. 32, 8940–8951.

19. Iwata, R., Casimir, P., Erkol, E., Boubakar, L., Planque, M., Gallego López, I.M., Ditkowska, M., Gaspariunaite, V., Beckers, S., Remans, D., et al. (2023). Mitochondria metabolism sets the species-specific tempo of neuronal development. Science 379, eabn4705.

20. Ji, S., Zhou, W., Li, X., Liu, S., Wang, F., Li, X., Zhao, T., Ji, G., Du, J., and Hao, A. (2019). Maternal hyperglycemia disturbs neocortical neurogenesis via epigenetic regulation in C57BL/6J mice. Cell Death Dis. 10, 211.

21. Fu, J., Tay, S.S.W., Ling, E.A., and Dheen, S.T. (2006). High glucose alters the expression of genes involved in proliferation and cell-fate specification of embryonic neural stem cells. Diabetologia 49, 1027–1038.

22. Rash, B.G., Micali, N., Huttner, A.J., Morozov, Y.M., Horvath, T.L., and Rakic, P. (2018). Metabolic regulation and glucose sensitivity of cortical radial glial cells. Proc. Natl. Acad. Sci. U. S. A. 115, 10142–10147.

23. Zheng, X., Boyer, L., Jin, M., Mertens, J., Kim, Y., Ma, L., Ma, L., Hamm, M., Gage, F.H., and Hunter, T. (2016). Metabolic reprogramming during neuronal differentiation from aerobic glycolysis to neuronal oxidative phosphorylation. Elife 5. 10.7554/eLife.13374.

24. Andrews, M.G., and Pearson, C.A. (2024). Toward an understanding of glucose metabolism in radial glial biology and brain development. Life Sci. Alliance 7. 10.26508/lsa.202302193.

25. Lange, C., Turrero Garcia, M., Decimo, I., Bifari, F., Eelen, G., Quaegebeur, A., Boon, R., Zhao, H., Boeckx, B., Chang, J., et al. (2016). Relief of hypoxia by angiogenesis promotes neural stem cell differentiation by targeting glycolysis. EMBO J. 35, 924–941.

26. Javaherian, A., and Kriegstein, A. (2009). A stem cell niche for intermediate progenitor cells of the embryonic cortex. Cereb. Cortex 19 *Suppl 1*, i70–i77.

27. Komabayashi-Suzuki, M., Yamanishi, E., Watanabe, C., Okamura, M., Tabata, H., Iwai, R., Ajioka, I., Matsushita, J., Kidoya, H., Takakura, N., et al. (2019). Spatiotemporally dependent vascularization is differently utilized among neural progenitor subtypes during neocortical development. Cell Rep. 29, 1113–1129.e5.

28. Pașca, A.M., Park, J.-Y., Shin, H.-W., Qi, Q., Revah, O., Krasnoff, R., O’Hara, R., Willsey, A.J., Palmer, T.D., and Pașca, S.P. (2019). Human 3D cellular model of hypoxic brain injury of prematurity. Nat. Med. 25, 784–791.

29. Bhaduri, A., Andrews, M.G., Mancia Leon, W., Jung, D., Shin, D., Allen, D., Jung, D., Schmunk, G., Haeussler, M., Salma, J., et al. (2020). Cell stress in cortical organoids impairs molecular subtype specification. Nature 578, 142–148.

30. Uzquiano, A., Kedaigle, A.J., Pigoni, M., Paulsen, B., Adiconis, X., Kim, K., Faits, T., Nagaraja, S., Antón-Bolaños, N., Gerhardinger, C., et al. (2022). Proper acquisition of cell class identity in organoids allows definition of fate specification programs of the human cerebral cortex. Cell 185, 3770–3788.e27.

31. Vértesy, Á., Eichmüller, O.L., Naas, J., Novatchkova, M., Esk, C., Balmaña, M., Ladstaetter, S., Bock, C., von Haeseler, A., and Knoblich, J.A. (2022). Gruffi: an algorithm for computational removal of stressed cells from brain organoid transcriptomic datasets. EMBO J., e111118.

32. Kadoshima, T., Sakaguchi, H., Nakano, T., Soen, M., Ando, S., Eiraku, M., and Sasai, Y. (2013). Self-organization of axial polarity, inside-out layer pattern, and species-specific progenitor dynamics in human ES cell-derived neocortex. Proc. Natl. Acad. Sci. U. S. A. 110, 20284–20289.

33. Andrews, M.G., Siebert, C., Wang, L., White, M.L., Ross, J., Morales, R., Donnay, M., Bamfonga, G., Mukhtar, T., McKinney, A.A., et al. (2023). LIF signaling regulates outer radial glial to interneuron fate during human cortical development. Cell Stem Cell. 10.1016/j.stem.2023.08.009.

34. Oliveira, F.R., Barros, E.G., and Magalhães, J.A. (2002). Biochemical profile of amniotic fluid for the assessment of fetal and renal development. Braz. J. Med. Biol. Res. 35, 215–222.

35. Velika, B., Birkova, A., Dudic, R., Urdzik, P., and Marekova, M. (2018). Selected physicochemical properties of amniotic fluid according to week of pregnancy. Bratisl. Lek. Listy 119, 175–179.

36. Ozgu-Erdinc, A.S., Cavkaytar, S., Aktulay, A., Buyukkagnici, U., Erkaya, S., and Danisman, N. (2014). Mid-trimester maternal serum and amniotic fluid biomarkers for the prediction of preterm delivery and intrauterine growth retardation. J. Obstet. Gynaecol. Res. 40, 1540–1546.

37. Leen, W.G., Willemsen, M.A., Wevers, R.A., and Verbeek, M.M. (2012). Cerebrospinal fluid glucose and lactate: age-specific reference values and implications for clinical practice. PLoS One 7, e42745.

38. Mas-Bargues, C., Sanz-Ros, J., Román-Domínguez, A., Inglés, M., Gimeno-Mallench, L., El Alami, M., Viña-Almunia, J., Gambini, J., Viña, J., and Borrás, C. (2019). Relevance of oxygen concentration in stem cell culture for regenerative medicine. Int. J. Mol. Sci. 20, 1195.

39. Pollen, A.A., Nowakowski, T.J., Chen, J., Retallack, H., Sandoval-Espinosa, C., Nicholas, C.R., Shuga, J., Liu, S.J., Oldham, M.C., Diaz, A., et al. (2015). Molecular identity of human outer radial glia during cortical development. Cell 163, 55–67.

40. Hansen, D.V., Lui, J.H., Parker, P.R.L., and Kriegstein, A.R. (2010). Neurogenic radial glia in the outer subventricular zone of human neocortex. Nature 464, 554–561.

41. Dehay, C., Kennedy, H., and Kosik, K.S. (2015). The outer subventricular zone and primate-specific cortical complexification. Neuron 85, 683–694.

42. Kalebic, N., Gilardi, C., Stepien, B., Wilsch-Bräuninger, M., Long, K.R., Namba, T., Florio, M., Langen, B., Lombardot, B., Shevchenko, A., et al. (2019). Neocortical Expansion Due to Increased Proliferation of Basal Progenitors Is Linked to Changes in Their Morphology. Cell Stem Cell 24, 535–550.e9.

43. Huang, Y., Mohanty, V., Dede, M., Tsai, K., Daher, M., Li, L., Rezvani, K., and Chen, K. (2023). Characterizing cancer metabolism from bulk and single-cell RNA-seq data using METAFlux. Nat. Commun. 14, 4883.

44. Demarquoy, J., and Le Borgne, F. (2015). Crosstalk between mitochondria and peroxisomes. World J. Biol. Chem. 6, 301–309.

45. Iwata, R., Gallego-Lopez, I.M., Erkol, E., Limame, R., Vandekeere, A., Benac, N., Turner-Bridger, B., Planque, M., Ditkowska, M., Lein, V., et al. (2025). Species-specific rates of fatty acid metabolism set the scale of temporal patterning of corticogenesis through protein acetylation dynamics. bioRxiv. 10.1101/2025.08.27.672586.

46. Andrews, M.G., Subramanian, L., and Kriegstein, A.R. (2020). mTOR signaling regulates the morphology and migration of outer radial glia in developing human cortex. Elife 9. 10.7554/eLife.58737.

47. Wang, Y., Chiola, S., Yang, G., Russell, C., Armstrong, C.J., Wu, Y., Spampanato, J., Tarboton, P., Ullah, H.M.A., Edgar, N.U., et al. (2022). Modeling human telencephalic development and autism-associated SHANK3 deficiency using organoids generated from single neural rosettes. Nat. Commun. 13, 5688.

48. Amelio, I., Cutruzzolá, F., Antonov, A., Agostini, M., and Melino, G. (2014). Serine and glycine metabolism in cancer. Trends Biochem. Sci. 39, 191–198.

49. Agostini, M., Romeo, F., Inoue, S., Niklison-Chirou, M.V., Elia, A.J., Dinsdale, D., Morone, N., Knight, R.A., Mak, T.W., and Melino, G. (2016). Metabolic reprogramming during neuronal differentiation. Cell Death Differ. 23, 1502–1514.

50. Dienel, G.A. (2019). Brain glucose metabolism: Integration of energetics with function. Physiol. Rev. 99, 949–1045.

51. Mil, J., Soto, J.A., Matulionis, N., Krall, A., Day, F., Stiles, L., Montales, K.P., Azizad, D.J., Gonzalez, C.E., Nano, P.R., et al. (2025). Metabolic atlas of early human cortex identifies regulators of cell fate transitions. bioRxivorg. 10.1101/2025.03.10.642470.

52. Florio, M., Albert, M., Taverna, E., Namba, T., Brandl, H., Lewitus, E., Haffner, C., Sykes, A., Wong, F.K., Peters, J., et al. (2015). Human-specific gene ARHGAP11B promotes basal progenitor amplification and neocortex expansion. Science 347, 1465–1470.

53. Namba, T., Dóczi, J., Pinson, A., Xing, L., Kalebic, N., Wilsch-Bräuninger, M., Long, K.R., Vaid, S., Lauer, J., Bogdanova, A., et al. (2020). Human-specific ARHGAP11B acts in mitochondria to expand neocortical progenitors by glutaminolysis. Neuron 105, 867–881.e9.

54. Kantarci, H., Gou, Y., and Riley, B.B. (2020). The Warburg Effect and lactate signaling augment Fgf-MAPK to promote sensory-neural development in the otic vesicle. Elife 9. 10.7554/eLife.56301.

55. Certo, M., Llibre, A., Lee, W., and Mauro, C. (2022). Understanding lactate sensing and signalling. Trends Endocrinol. Metab. 33, 722–735.

56. Mu, X., Shi, W., Xu, Y., Xu, C., Zhao, T., Geng, B., Yang, J., Pan, J., Hu, S., Zhang, C., et al. (2018). Tumor-derived lactate induces M2 macrophage polarization via the activation of the ERK/STAT3 signaling pathway in breast cancer. Cell Cycle 17, 428–438.

57. Ostrem, B.E.L., Lui, J.H., Gertz, C.C., and Kriegstein, A.R. (2014). Control of outer radial glial stem cell mitosis in the human brain. Cell Rep. 8, 656–664.

58. Ma, H., Folmes, C.D.L., Wu, J., Morey, R., Mora-Castilla, S., Ocampo, A., Ma, L., Poulton, J., Wang, X., Ahmed, R., et al. (2015). Metabolic rescue in pluripotent cells from patients with mtDNA disease. Nature 524, 234–238.

59. Romero-Morales, A.I., Robertson, G.L., Rastogi, A., Rasmussen, M.L., Temuri, H., McElroy, G.S., Chakrabarty, R.P., Hsu, L., Almonacid, P.M., Millis, B.A., et al. (2022). Human iPSC-derived cerebral organoids model features of Leigh syndrome and reveal abnormal corticogenesis. Development 149. 10.1242/dev.199914.

60. Pinto Payares, D.V., Spooner, L., Vosters, J., Dominguez, S., Patrick, L., Harris, A., and Kanungo, S. (2024). A systematic review on the role of mitochondrial dysfunction/disorders in neurodevelopmental disorders and psychiatric/behavioral disorders. Front. Psychiatry 15, 1389093.

61. Brister, D., Werner, B.A., Gideon, G., McCarty, P.J., Lane, A., Burrows, B.T., McLees, S., Adelson, P.D., Arango, J.I., Marsh, W., et al. (2022). Central nervous system metabolism in autism, epilepsy and developmental delays: A cerebrospinal fluid analysis. Metabolites 12, 371.

62. Brister, D., Rose, S., Delhey, L., Tippett, M., Jin, Y., Gu, H., and Frye, R.E. (2022). Metabolomic signatures of autism Spectrum Disorder. J. Pers. Med. 12, 1727.

63. Koch, H., and Weber, Y.G. (2019). The glucose transporter type 1 (Glut1) syndromes. Epilepsy Behav. 91, 90–93.

64. Matsuda, T., and Cepko, C.L. (2004). Electroporation and RNA interference in the rodent retina in vivo and in vitro. Proc. Natl. Acad. Sci. U. S. A. 101, 16–22.

65. Fellmann, C., Hoffmann, T., Sridhar, V., Hopfgartner, B., Muhar, M., Roth, M., Lai, D.Y., Barbosa, I.A.M., Kwon, J.S., Guan, Y., et al. (2013). An optimized microRNA backbone for effective single-copy RNAi. Cell Rep. 5, 1704–1713.

66. 66. Pollen, A.A., Bhaduri, A., Andrews, M.G., Nowakowski, T.J., Meyerson, O.S., Mostajo-Radji, M.A., Di Lullo, E., Alvarado, B., Bedolli, M., Dougherty, M.L., et al. (2019). Establishing Cerebral Organoids as Models of Human-Specific Brain Evolution. Cell 176, 743–756.e17.

67. Gu, H., Zhang, P., Zhu, J., and Raftery, D. (2015). Globally optimized targeted mass spectrometry: Reliable metabolomics analysis with broad coverage. Anal. Chem. 87, 12355–12362.

68. Wei, Y., Jasbi, P., Shi, X., Turner, C., Hrovat, J., Liu, L., Rabena, Y., Porter, P., and Gu, H. (2021). Early breast cancer detection using untargeted and targeted metabolomics. J. Proteome Res. 20, 3124–3133.

69. Jin, Y., Chi, J., LoMonaco, K., Boon, A., and Gu, H. (2023). Recent review on selected xenobiotics and their impacts on gut microbiome and metabolome. Trends Analyt. Chem. 166. 10.1016/j.trac.2023.117155.

70. Garcia, M.M., Romero, A.S., Merkley, S.D., Meyer-Hagen, J.L., Forbes, C., Hayek, E.E., Sciezka, D.P., Templeton, R., Gonzalez-Estrella, J., Jin, Y., et al. (2024). In vivo tissue distribution of polystyrene or mixed polymer microspheres and metabolomic analysis after oral exposure in mice. Environ. Health Perspect. 132, 47005.

71. Scieszka, D.P., Garland, D., Hunter, R., Herbert, G., Lucas, S., Jin, Y., Gu, H., Campen, M.J., and Cannon, J.L. (2023). Multi-omic assessment shows dysregulation of pulmonary and systemic immunity to e-cigarette exposure. Respir. Res. 24, 138.

72. Kim, S., Li, H., Jin, Y., Armad, J., Gu, H., Mani, S., and Cui, J.Y. (2023). Maternal PBDE exposure disrupts gut microbiome and promotes hepatic proinflammatory signaling in humanized PXR-transgenic mouse offspring over time. Toxicol. Sci. 194, 209–225.

73. Mohr, A.E., Sweazea, K.L., Bowes, D.A., Jasbi, P., Whisner, C.M., Sears, D.D., Krajmalnik-Brown, R., Jin, Y., Gu, H., Klein-Seetharaman, J., et al. (2024). Gut microbiome remodeling and metabolomic profile improves in response to protein pacing with intermittent fasting versus continuous caloric restriction. Nat. Commun. 15, 4155.

74. Pelossof, R., Fairchild, L., Huang, C.-H., Widmer, C., Sreedharan, V.T., Sinha, N., Lai, D.-Y., Guan, Y., Premsrirut, P.K., Tschaharganeh, D.F., et al. (2017). Prediction of potent shRNAs with a sequential classification algorithm. Nat. Biotechnol. 35, 350–353.

75. Stirling, D.R., Swain-Bowden, M.J., Lucas, A.M., Carpenter, A.E., Cimini, B.A., and Goodman, A. (2021). CellProfiler 4: improvements in speed, utility and usability. BMC Bioinformatics 22, 433.

76. Dieterle, F., Ross, A., Schlotterbeck, G., and Senn, H. (2006). Probabilistic quotient normalization as robust method to account for dilution of complex biological mixtures. Application in 1H NMR metabonomics. Anal. Chem. 78, 4281–4290.

77. MetaboAnalyst https://www.metaboanalyst.ca/.

78. Hao, Y., Stuart, T., Kowalski, M.H., Choudhary, S., Hoffman, P., Hartman, A., Srivastava, A., Molla, G., Madad, S., Fernandez-Granda, C., et al. (2024). Dictionary learning for integrative, multimodal and scalable single-cell analysis. Nat. Biotechnol. 42, 293–304.

79. Choudhary, S., and Satija, R. (2022). Comparison and evaluation of statistical error models for scRNA-seq. Genome Biol. 23, 27.

80. Hao, Y., Hao, S., Andersen-Nissen, E., Mauck, W.M., 3rd, Zheng, S., Butler, A., Lee, M.J., Wilk, A.J., Darby, C., Zager, M., et al. (2021). Integrated analysis of multimodal single-cell data. Cell 184, 3573–3587.e29.

81. Love, M.I., Huber, W., and Anders, S. (2014). Moderated estimation of fold change and dispersion for RNA-seq data with DESeq2. Genome Biol. 15, 550.

82. Blighe, K. (2018). EnhancedVolcano (Bioconductor) 10.18129/B9.BIOC.ENHANCEDVOLCANO.

83. Yu, G., Wang, L.-G., Han, Y., and He, Q.-Y. (2012). clusterProfiler: an R package for comparing biological themes among gene clusters. OMICS 16, 284–287.

